# Interplay of pericentromeric genome organization and chromatin landscape regulates the expression of *Drosophila melanogaster* heterochromatic genes

**DOI:** 10.1101/534065

**Authors:** Parna Saha, Divya Tej Sowpati, Ishanee Srivastava, Rakesh Kumar Mishra

## Abstract

Transcription of heterochromatic genes residing within the constitutive heterochromatin is paradoxical to the tenets of the epigenetic code. *Drosophila melanogaster* heterochromatic genes serve as an excellent model system to understand the mechanisms of their transcriptional regulation. Recent developments in chromatin conformation techniques have revealed that genome organization regulates the transcriptional outputs. Thus, using 5C-seq in S2 cells, we present a detailed characterization of the hierarchical genome organization of *Drosophila* pericentromeric heterochromatin and its contribution to heterochromatic gene expression. We show that pericentromeric TAD borders are enriched in nuclear Matrix attachment regions while the intra-TAD interactions are mediated by various insulator binding proteins. Heterochromatic genes of similar expression levels cluster into Het TADs which indicates their transcriptional co-regulation. To elucidate how heterochromatic factors, influence the expression of heterochromatic genes, we performed 5C-seq in the HP1a or Su(var)3-9 depleted cells. HP1a or Su(var)3-9 RNAi results in perturbation of global pericentromeric TAD organization but the expression of the heterochromatic genes is minimally affected. Subset of active heterochromatic genes have been shown to have combination of HP1a/H3K9me3 with H3K36me3 at their exons. Interestingly, the knock-down of dMES-4 (H3K36 methyltransferase), downregulates expression of the heterochromatic genes. This indicates that the local chromatin interactions and the combination of heterochromatic factors (HP1a or H3K9me3) along with the H3K36me3 is crucial to drive the expression of heterochromatic genes. Furthermore, dADD1, present near the TSS of the active heterochromatic genes, can bind to both H3K9me3 or HP1a and facilitate the heterochromatic gene expression by regulating the H3K36me3 levels. Therefore, our findings provide mechanistic insights into the interplay of genome organization and chromatin factors at the pericentromeric heterochromatin that regulates *Drosophila melanogaster* heterochromatic gene expression.

## Introduction

Expression of genes present in the transcriptionally-inert heterochromatin is counter-intuitive and their mechanisms of expression remain elusive. Such genes are called the heterochromatic genes due to their location in the pericentromeric regions and their dependence on heterochromatic *trans* factors for their expression. Examples across species, include the pericentromeric genes of *Drosophila melanogaster* (Dimitri et al. 2003, 2009), juxtacentromeric genes (Beckers et al. 2001; Ruault et al. 2002; Brun et al. 2011) and X-chromosome inactivation escapee genes (Balaton and Brown 2016) in mammals and the genes in the centromeric knob regions in *Arabidopsis* (Bennetzen 2000; Le et al. 2015). *Drosophila melanogaster* heterochromatic genes are one of the most well-studied (N Corradini et al. 2007) owing to the strength of genetic manipulations that made interrogating the heterochromatic sequences possible even before the advent of the high-throughput sequencing technologies. They were identified by complementation analyses using chromosomal rearrangements (Hilliker 1976) and later by mapping the genes using fluorescence *in-situ* hybridization to the pericentromeric regions (Nicoletta Corradini et al. 2003). These heterochromatic genes encode proteins involved in various cellular processes, for example-*light*–transporter protein; *rolled* – MAP kinase involved in imaginal disc development; *concertina*-gastrulation protein; *NippedA, NippedB*– transcription regulators and ribosomal proteins like *Rpl 5/15/38* and many more which still remain uncharacterized. The comparative studies on the evolution of the heterochromatic genes in the Drosophilids, suggest that the promoter structure of the *Drosophila melanogaster* heterochromatic genes is indistinguishable from the euchromatic orthologues in the older *Drosophila* species like *D.virilis and D.psuedoobscura*. The large introns of the *D melanogaster* heterochromatic genes have an accumulation of transposable elements and thus these genes have been pushed into heterochromatin. However, they have adapted to stay active while keeping the intronic TEs silent, using regulatory mechanisms that are largely unknown (Dimitri, Junakovic, and Arcà 2003; Yasuhara, DeCrease, and Wakimoto 2005). The reports of abrogation of heterochromatic gene expression upon chromosomal translocation of these genes into the euchromatin (Wakimoto and Hearn 1990; Yasuhara and Wakimoto 2008; Clegg et al. 1998) or the depletion of heterochromatic factors (Eberl, Duyf, and Hilliker 1993; Lu et al. 2000) further indicated the need of heterochromatic environment for their expression. Thus, the expression of heterochromatic genes is contradictory to the tenets of heterochromatin and there exists a limited understanding of how the pericentromeric genome organization and epigenetic landscape contribute to their transcription.

The earliest evidence of the contribution of the genomic context in determining heterochromatin gene regulation was indicated by the report of reciprocal PEV for the heterochromatic genes (Schultz 1936; Hessler 1957). Subsequently, it was further reported that the extent of variegation of the *lt (light)* gene is correlated to the site of heterochromatic breakpoint-centromere-proximal heterochromatic genes were almost unaffected while the centromere distal heterochromatic gene showed the most severe phenotypes upon translocation (Wakimoto and Hearn 1990). Thus, it has been believed that the genomic organization of the heterochromatic genes and the local concentration of certain heterochromatic *trans* factors is deterministic for their expression. Furthermore, the epigenetic landscape of the heterochromatic genes is rather unique with the combinatorial occurrence of active and inactive histone modifications like H3K4me3/1, H3K9/27/14 ac at the TSS and H3K9me3, H3K36me3 and HP1 on the gene body (Nicole C. Riddle et al. 2011; Saha, Sowpati, and Mishra 2018). These interesting and unique combinations of the histone marks are thought to be responsible for marking these genes for transcription, differentially from the surrounding repeat-rich repressive heterochromatin. With the recent surge in the genome organization studies, it is being increasingly appreciated that structural partitioning of the genome into epigenetic domains and the transcriptional outputs are inter-related (Wit and Laat 2012; Yu and Ren 2017). Despite the fact that the *Drosophila* genome organization into various chromatin domains and its relationship to gene expression in various developmental contexts has been well-studied (Schwartz and Cavalli 2017), the interplay of genome organization in the pericentromeric heterochromatin and heterochromatic proteins in regulating heterochromatic gene expression remains largely uninvestigated.

In this study, we sought to dissect the inter-dependence of pericentromeric epigenetic landscape and genome organization in heterochromatic gene regulation. We present the first report of the long-range DNA interactome at the pericentromeric heterochromatin and characterized those pericentromere associated domains (Het TADs) with respect to several genomic and epigenomic features. We report that this structural organization correlates with the expression patterns of the heterochromatic genes. Furthermore, we also delineated the role of two crucial heterochromatin protein (HP1a and Su(var)3-9) at the level of pericentromeric genome organization and also, the heterochromatic gene expression. Additionally, we investigated the roles of two chromatin proteins-dMES-4 and dADD1 in regulating heterochromatic gene expression, whose contribution has not been reported earlier. Despite having clues about a few candidate heterochromatic genes, a well-validated model of heterochromatic gene expression in *Drosophila* has been lacking. Our study has been focused on filling this lacuna and gaining a global understanding of this intriguing biological phenomenon. Taken together our findings, show that the local long-range DNA interactions in the pericentromeres are sufficient to maintain the heterochromatic gene expression in the presence of transcriptionally favourable chromatin landscape.

## Results

### 1. Pericentromeric heterochromatin is organised into distinct TADs

To investigate the pericentromeric genome organization and its contribution to heterochromatic gene expression, we sought to map the long-range interactions in the pericentromeric regions of the *Drosophila melanogaster* genome. Exhaustive information of the chromatin contacts in the heterochromatin has been lacking (Sexton et al. 2012) owing to the discrepancy in mapping repetitive sequences and the existing Hi-C data lacks sufficient information about these regions (Supplementary Figure S1) as seen using Chorogenome Navigator (Ramírez et al. 2018). Thus, we performed chromosome conformation capture carbon copy or 5C (Dostie and Dekker 2007) targeted at the pericentromeric regions as per the annotation of the euchromatic-heterochromatic junctions in the dm3 genome build (Smith et al. 2007; Hoskins et al. 2007), **Figure 1A**. We also included 100kb of the adjacent euchromatic regions on each chromosome arm. 678 primes were designed at the EcoRI sites to interrogate the pericentromeric regions on Chr2L, Chr2R, Chr3L, Chr3R and ChrX that covered 6Mb of the *Drosophila melanogaster* genome and approximately 250 heterochromatic genes. The 3C and 5C libraries were prepared from *Drosophila melanogaster* S2 cells and proper quality controls were ensured (Supplementary Figure S2-A-B). We selected paired-end reads with the EcoRI site which were filtered to remove the low-quality reads. Using the my5C tools, we mapped 3540 interactions on Chr2L, 4900 interactions on Chr2R, 6142 interactions on Chr3L, 900 interactions on Chr3R and 1680 interactions on ChrX respectively, that are plotted as heatmaps based on their interaction frequency (Supplementary Figure S3). A closer inspection of the intra chromosomal interactions reveal that there are several local domains of interacting regions along the pericentromeric arm distinct from the self-interacting regions in the euchromatin (**Figure 1B**). This indicates that the pericentromeric regions partition into domains distinct from the euchromatic regions. We also mapped several inter-chromosomal interactions amongst the Het regions ((Supplementary Figure S3-B), which are likely to be functionally rellevant given that the centromeres coalesce into chromocenter in the *Drosophila* nuclei. Next, we proceeded to computationally define the TADs and sub TADs using the Directionality index scores based on the interactions mapped in each of the 3 biological replicates across the 5 chromosome arms (Supplementary Figure S4-A). The correlation between the replicates was achieved using Hidden Markov Model (HMM) to correct biases and the TAD border span was decided upon using average insulated region (Figure S3-B). We report 23 TADs (5 on Chr2L, 7 on Chr2R, 6 on chr3L, 5 on ChrX) with the average TAD sizes of 50-100kb and TAD border sizes in the range of 20-40kb (Table 1 and **Figure 1B**). The interactions mapped to Chr 3R were not enough to demarcate TAD computationally. The interactions reported in the 5C experiments were further validated using 3C PCR (Supplementary Figure S5-C). We also validated the interactions in the Chr2L involving regions of actively expressed genes coming together using fluorescence *in-situ* where two interacting regions are 280kb apart on the linear scale (Supplementary Figure S5-A-B). The repertoire of long-range DNA interactions obtained, provides the first report of the presence of distinct higher order genomic architecture within the condensed *Drosophila melanogaster* chromocenter.

**Figure 1:**
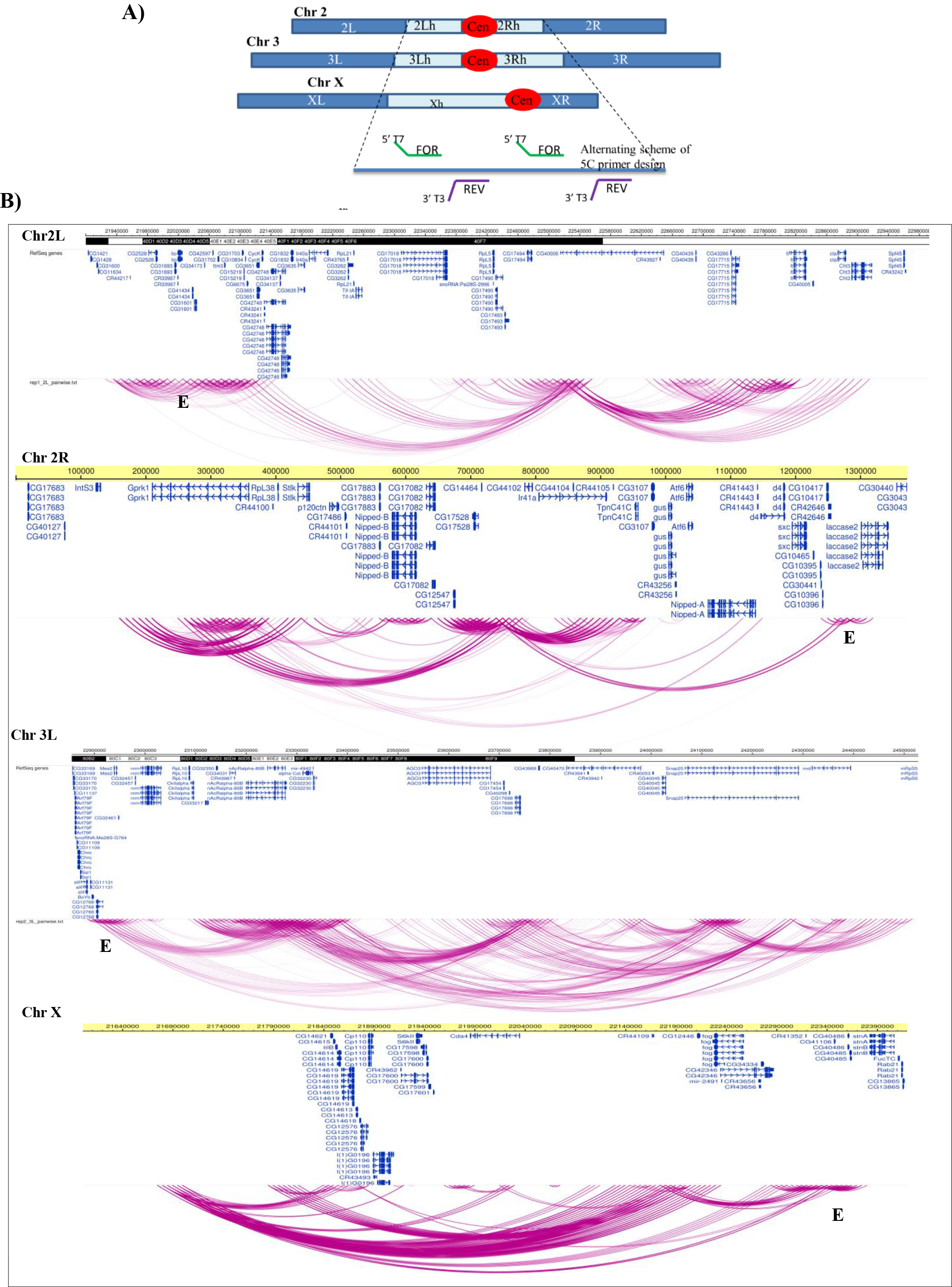
Pericentromeric heterochromatin is organized into domains. **A)** Representation of the *Drosophila melanogaster* chromosomes where the pericentromeric regions are marked in light blue are the regions included in the 5C experiment. Cen in red oval denotes the centromere. Euchromatin is shown in dark blue in each arm. Zoomed in schematic representation depicts 5C primer design by alternating primer design scheme, where forward (with T7 overhang) and reverse (with T3 overhang) primers are designed at the alternate restriction enzyme (EcoRI) cut site. **B)** Representation of the 5C interaction domains and subdomains mapped the pericentromeric regions of Chr 2L, Chr 2R, Chr 3L, Chr X along with the Refseq genes, using Washington University epigenome browser. Euchromatic regions included in the study partitions into distinct domains (E).

### 2. Characteristics of the *Drosophila* Het TADs

In order to characterize the TADs present in the pericentromeric heterochromatin (Het TADs), we overlapped the intra-TAD interactions and TAD borders identified in our 5C experiment with various genomic and epigenomic features. This included a) genomic features – enhancers (STARR seq), Nuclear **M**atrix **A**ssociated **R**egions (MARs), mapped in *Drosophila* S2 cells (unpublished data from lab), predicted boundary elements (BEs) from the cdBEST tool (Srinivasan and Mishra 2012) and various classes of repeats; b) enrichment of various insulator binding proteins (IBPs) like dCTCF, BEAF32, GAF, CP190, Su(Hw), mod(mdg4) and c) epigenomic features like histone modifications and heterochromatin proteins (HP1a/b/c). To ascribe statistical significance to the observations, randomized regions were overlapped with the features and p-value was assigned as a part of the OverlapPerm test for TAD borders with each of them (Supplementary Figure S6 and S7). The overlap of TAD borders with MARs was highly significant (p-value 0.001), **Figure 2A**.

**Figure 2:**
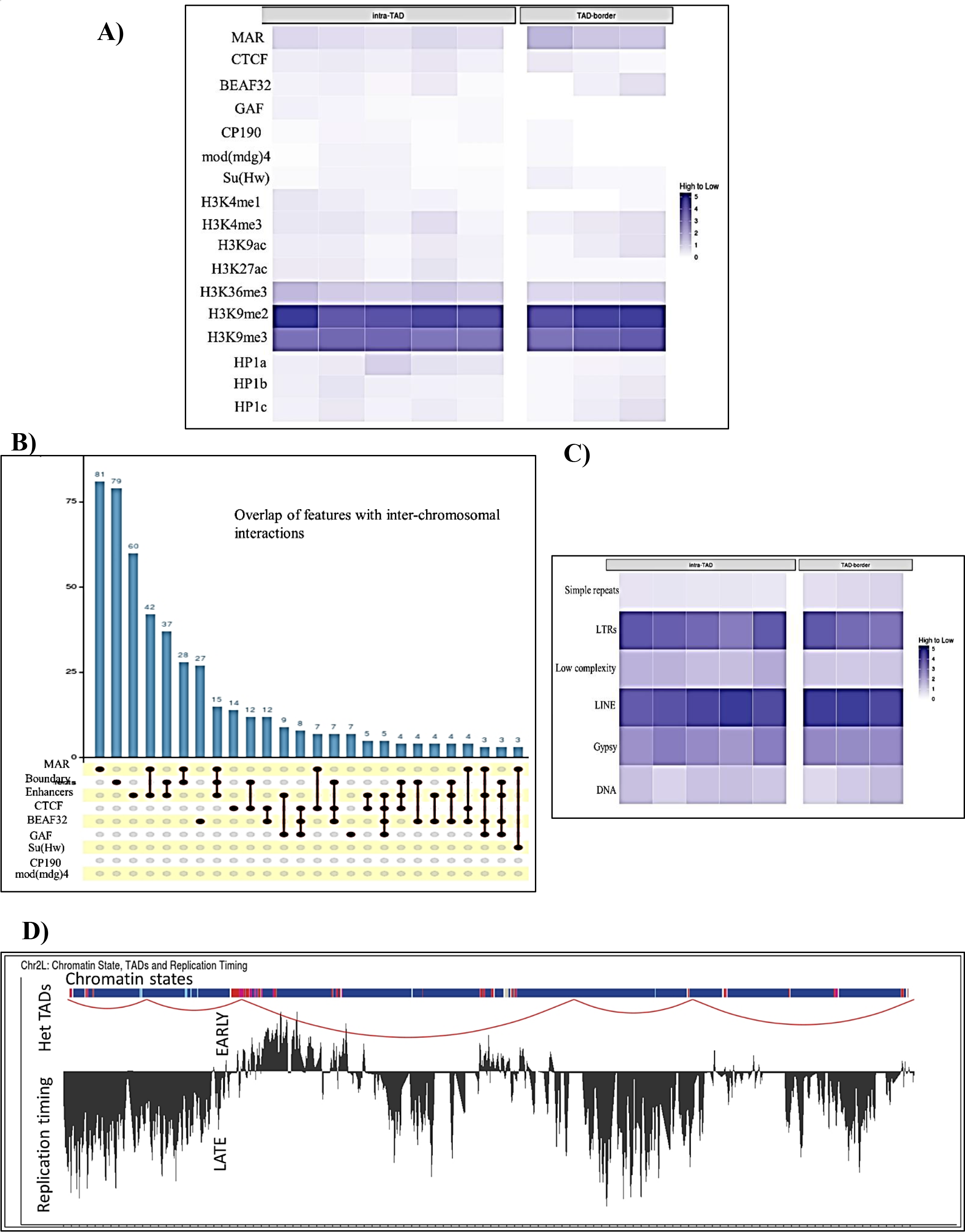
Characteristics of the pericentromeric Het TADs. **A)** Comparative heatmap for the enrichment of various genomic and epigenomic features across Het TAD borders and intra TAD interactions **-** TAD borders are enriched in MARs. Inter TAD interactions overlap enriched for BEAF32 and dCTCF peaks. GAF present predominantly at the intra TAD interactions. Among histone modifications, the active marks like H3K4me1, H3K4me3, H3K9ac are present within the TADs whereas H3K27ac is depleted at the TAD borders. H3K36me3 is present both at the intra TAD and TAD border regions as are the heterochromatic marks H3K9me2/3. HP1a, HP1b and HP1c are more enriched within the TAD regions. **B)** Characteristics of inter-chromosomal interactions with respect to various genomic feature - MARs, enhancers, cdBEST boundary elements and ChIP peaks of insulator proteins BEAF32, GAF, CTCF, CP190, Su(Hw), mod(mdg)4. The y-axis indicates the number of unique overlaps mentioned on the top of the bar graph with a particular feature (black dots) or combinations of feature (denoted by connected black dots). **C)** Overlap of the interaction data with various repeat elements shows no difference between the enrichment at the intra TAD regions and TAD borders. **D)** TAD borders also mark replication timing domains - Representative image showing the overlap of Het TADs of Chr2L with the replication timing domains. TAD borders mark the domains of replication timing and are early replicating as compared to the intra TAD regions. Chromatin states are as per the 9 state chromatin model (Kharchenko *et al*; *Nature* 2011) where red depicts State-1 in red (active transcription and exons marked by H3K4me3/H3K9ac) and State-7 in blue (H3K9me3).

We find that the TAD borders are enriched in MARs, followed by cdBEST boundary elements. Enhancers are less abundant at the TAD borders (p-value 0.34) indicating that the short-range intra-TAD interactions are involved in enhancer-promoter contacts. We were also interested in characterizing the heterochromatic TADs in terms of binding of various architectural and insulator proteins which have been reported to mediate genome organization in flies and used the available ChIP datasets for the same (Supplementary Table 5). The relative enrichment heat–map for the intra TAD interactions and TAD borders, **Figure 2A**, shows that the TAD borders are enriched in dCTCF (p-value 0.002) and BEAF32 (p-value 0.001 (for Chr 3L), 0.002 (for ChrX)). While DmGAF (p-value 0.002), CP190 (p-value 0.012), Su(Hw) (p-value 0.136) and mod(mdg4) (p-value 0.001) Supplementary Figure S7) are present in the intra-TAD regions. dCTCF has no discernable difference in its enrichment at either TAD borders or the intra TAD regions. H3K36me3 and H3K9me2/3 are present both at the TAD borders and intra-TAD regions. Inter-chromosomal interactions are also enriched for MARs followed by various combinations of insulator binding proteins-BEAF32 and dCTCF being the predominant factors mediating the inter TAD interactions, **Figure 2B**.

Given, the heterochromatic regions are enriched in various transposable elements and repeats we attempted to understand whether they have any role in genome organization and are thus preferentially enriched at the TAD borders. We chose all available classes of repeat elements and overlapped with the TAD borders identified in our study. No difference in the preference for either the intra TAD regions or the TAD borders was noted, **Figure 2C**. Thus, there is no apparent functional correlation of repeat occurrence with the pericentromeric genome organization. We also overlapped the 5C interactions with 9 chromatin states and the replication timing data from S2 cells. We report that the Het TADs also mark the replication timing domains in the S2 cells and active Het TAD borders tend to overlap with the early-replicating regions, **Figure 2D** and Supplementary Figure S8. These observations confirm that the Het TAD boundaries are marked by MARs while the inter-TAD interactions by various combinations of insulator binding proteins and collectively this genomic organization likely to regulate important genomic functions (transcription, replication) in the heterochromatin.

### 3. Heterochromatic genes with similar expression levels in S2 cells are clustered together in Het TADs

TADs reported in *Drosophila melanogaster* has been shown to partition the genome into different epigenetic and thus expression domains (Sexton et al. 2012). Therefore, we asked whether the genomic organization observed at the pericentromeric region could be related to heterochromatic gene expression. To this end, we performed a total RNA seq on the S2 cells. We identified transcripts with a strong correlation between the 3 biological replicates (PCC >0.96). In order to verify our hypothesis, we overlapped the transcriptomic data with the Het TADs identified in our study. We find that across the heterochromatic regions of 5 chromosome arms, the TAD organization correlates with the presence of similarly expressing heterochromatic genes within them, **Figure 3** and Supplementary Figure S9. The Het TAD borders, similar to the euchromatic regions, are mostly enriched in genes having high to moderate expression in S2 cells, **Table 1**. This organization of the TADs divides the pericentromeric genome into patches of active and inactive TADs within the seemingly completely repressive heterochromatin, and is likely to mediate the transcriptional co-regulation of the heterochromatic genes.

**Figure 3:**
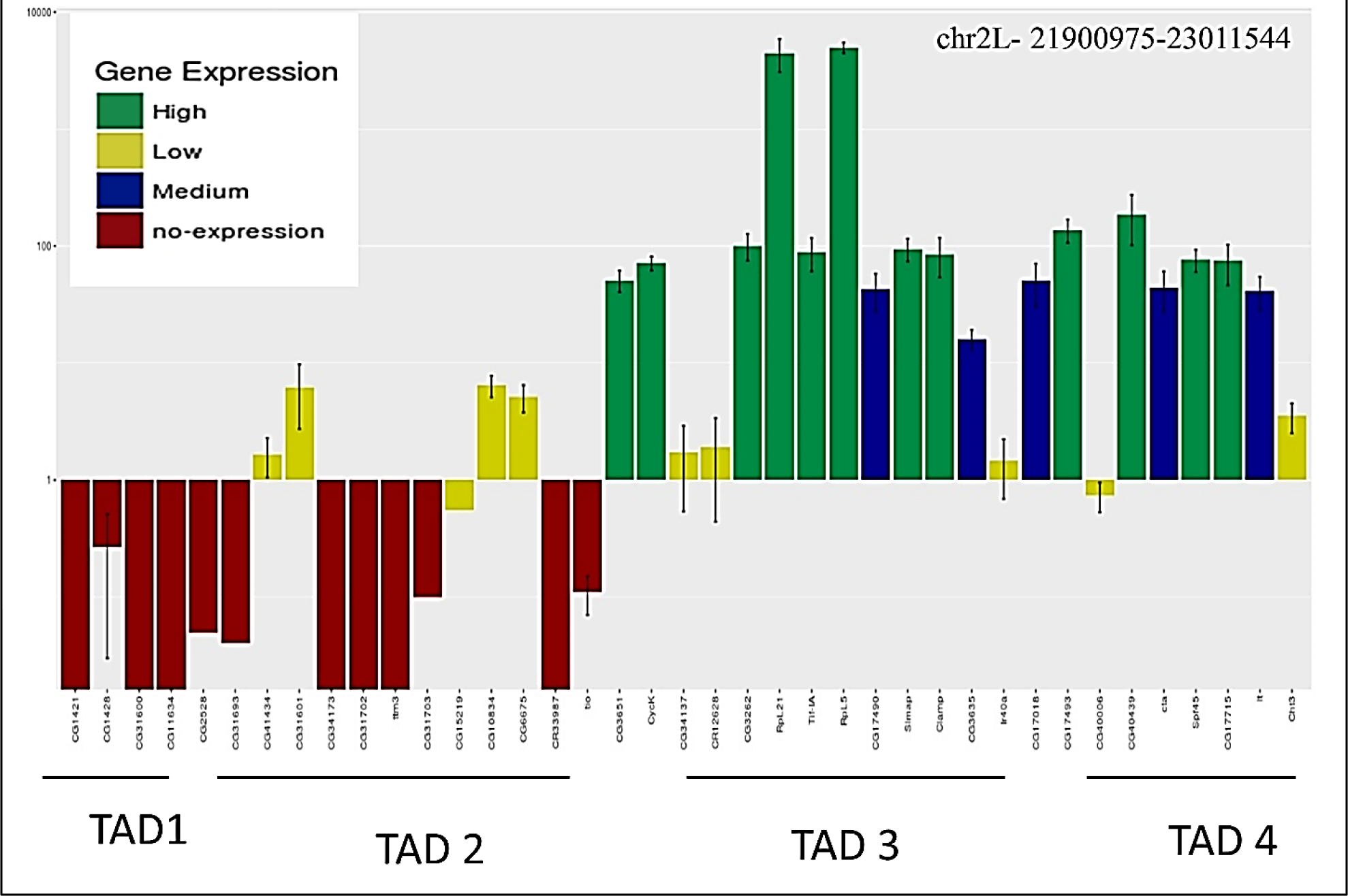
Correlation between pericentromeric genome organization and heterochromatic gene expression-. The representative image of Chr2L showing the overlap of RNA seq data with Het TADs. Bar plots showing the RNA seq data for each of the heterochromatic genes on the Chr2L pericentromere (n=3), colour-coded according to the expression levels. Heterochromatic genes of similar expression levels in the S2 cells are present in the same TAD indicating that the structural organization contributes to their transcriptional co-regulation.

### 4. Loss of heterochromatic dosage weakens Het TAD insulation but does not disrupt heterochromatic gene expression

Genetic studies using candidate based approach have shown that the heterochromatic genes depend on the heterochromatic environment for their expression (Eberl, Duyf, and Hilliker 1993; Lu et al. 2000). In flies, HP1a and Su(var)3-9 (the major H3K9methyl transferase) are the two major players involved in maintenance of the heterochromatin structure and function (Elgin and Reuter 2013). Therefore, we asked how would the genome organization and thus, heterochromatic gene expression change at the genome-wide level, upon depletion of these two proteins. We performed RNAi mediated knockdown of HP1a and Su(var)3-9 individually in two biological replicates. S2 cells transfected with dsRNA against GFP was used as a control. The knock-down of the proteins was validated on western blots using specific antibodies, **Figure 4A**. Following this, the HP1a or Su(var)3-9 RNAi treated S2 cells were used for generating 5C libraries. Upon comparison of the long-range interactions observed in HP1a or Su(var)3-9 RNAi depleted cells with the control, we observed that in both the knock-down conditions there are more interactions further away from the diagonal indicating inter TAD interactions and the loss of TAD insulation increase in both inter and intra TAD interactions, **Figure 4B** and Supplementary Figure S10-A. The increase in inter-TAD interactions was more pronounced in the case of HP1a RNAi, **Figure 4C** and also there is gain of new long-range interactions of high interaction score. We find that the number of intra-TAD interactions in Su(var)3-9 is less as compared to WT, indicating that HP1a and Su(var)3-9 have distinct roles in regulating heterochromatin organization. Overall, the absence of HP1a or Su(var)3-9 affects global TAD organization in the pericentromeric, albeit to different extents.

**Figure 4:**
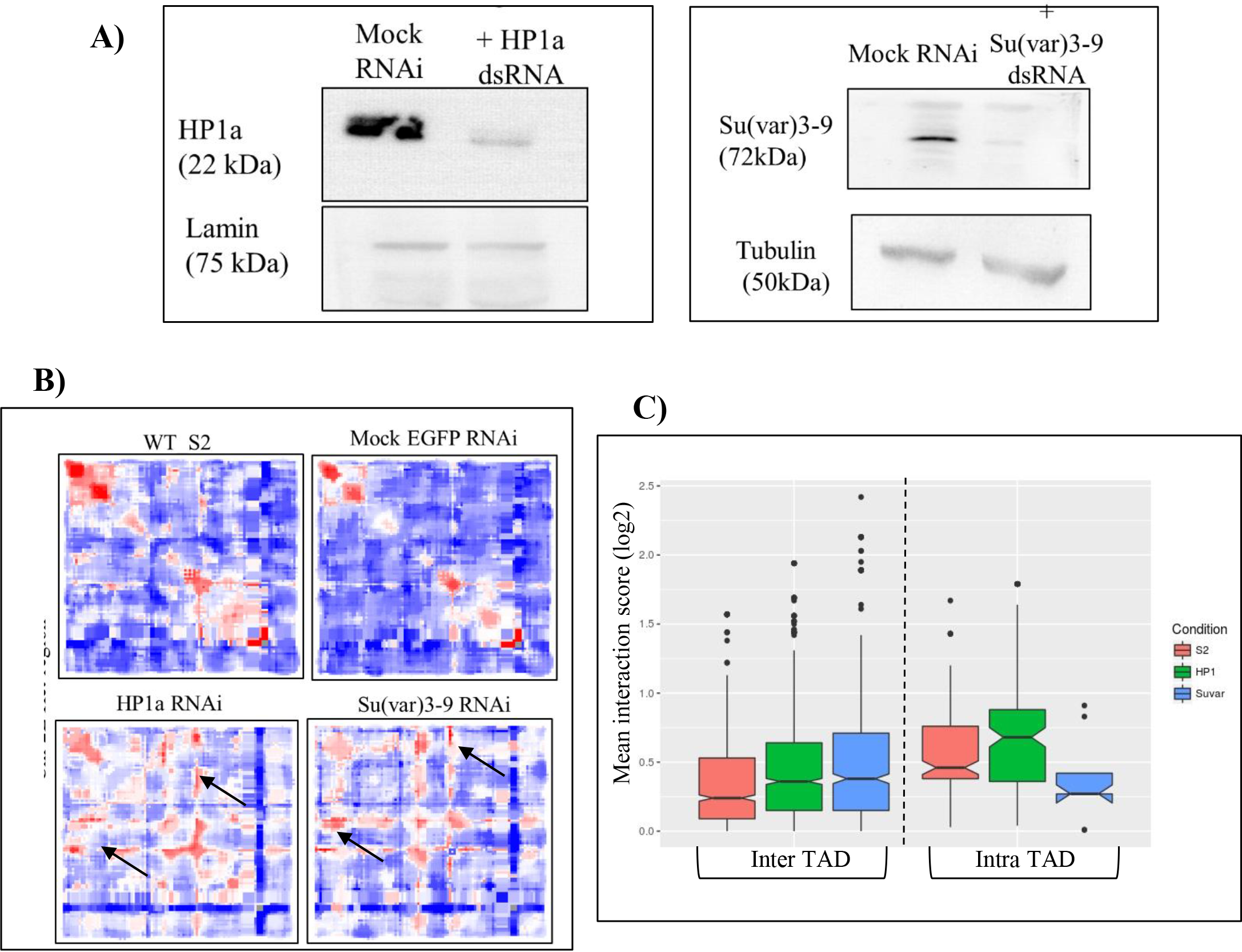
Depletion of HP1a and Su(var)3-9 by RNAi results in loss of TAD insulation. **A)** Knock-down of HP1a and Su(var)3-9 in S2 cells using dsRNA mediated RNA interference is confirmed by western blot. **B)** Comparison of the heat maps across the heterochromatic regions of Chr 2L included in the 5C seq in WT S2, mock RNAi, HP1a RNAi and Su(var)3-9 RNAi treated cells. The increase in inter-TAD interactions is evident by the increase in the number of red dots on either side of the diagonal of the heat map. **C)** Quantitation of the effect of knock-down of HP1a and Su(var)3-9 on the inter and intra TAD interactions using notched box plot where the notch indicates the median value at 95% confidence interval. The median notch of the WT and HP1a/ Su(var)3-9 RNAi conditions do not coincide indicating the gain of new interactions, some of which have high interaction scores and denoted as outliers.

Next, we asked how does the HP1a or Su(var)3-9 RNAi mediate disruption in TAD structure correlates with the expression of heterochromatic genes. To this end, we performed transcriptomic analyses on the HP1a or Su(var)3-9 RNAi treated cells. At the global level, a sizeable number of genes are affected in both HP1a and Su(var)3-9 RNAi conditions and 101 common genes are differentially regulated among the two RNAi conditions (Supplementary Figure S10-B). This is in line with the previous reports where both euchromatic and heterochromatic genes were shown to be affected by the depletion of heterochromatic proteins like HP1a. This indicates that HP1a and H3K9me3 marks have both activating and repressing effects depending upon the genomic context. Contrary to what was expected, we find down-regulation in the expression of only a few heterochromatic genes, **Figure 5A-D**. We observed that the heterochromatic genes that are affected (Supplementary Table 2), indicated by red stars (FDR < 0.05), are mostly located in the vicinity of the TAD borders. The heterochromatic genes affected by the Su(var)3-9 KD was lesser than those in case of HP1a. This prompted us to look into the levels of other methyltransferases like G9a and Setdb1 whose levels were uncompromised with Su(var)3-9 KD and therefore could have compensated for its function. Because the depletion of HP1a majorly causes an increase in the inter TAD interactions we believe that the local intra TAD interactions are still retained to facilitate the expression of the heterochromatic genes within the TADs and therefore the genes affected are mostly close to or at the TAD borders. This indicates that the global TAD organization at the pericentromeres is dispensable to maintain the expression of the heterochromatic genes.

**Figure 5:**
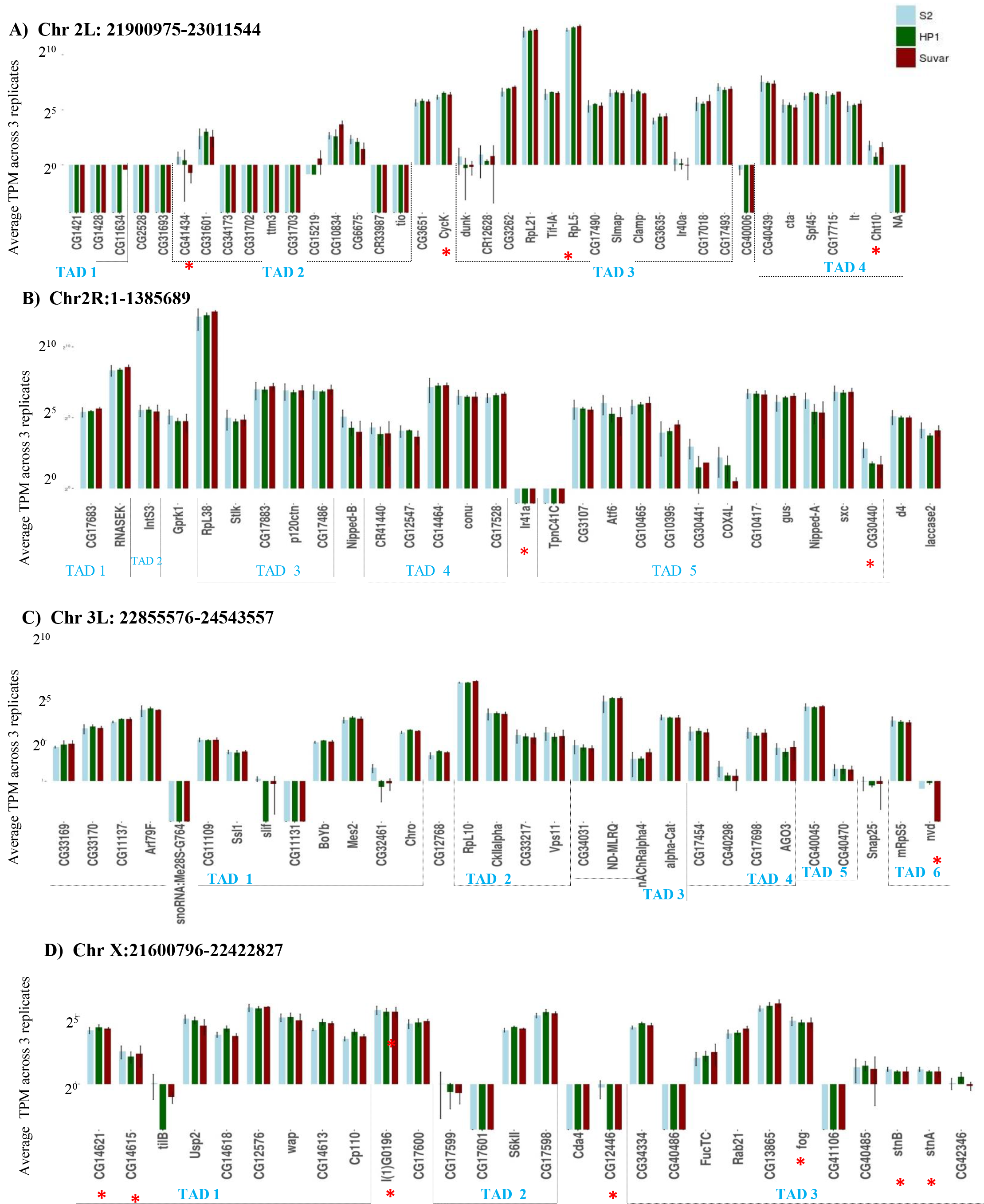
HP1a or Su(var) 3-9 KD has modest effect on the expression of heterochromatic genes-. Bar plot showing the effect of HP1a/Su(var)3-9 knock-down on the expression levels of heterochromatic genes in each of the Het TADs on each of the chromosome arm heterochromatin Chr 2L, **B)** Chr 2R, **C)** Chr3L and **D)** Chr X. Y-axis denotes the expression levels in log2 scale (n=3, p-value < 0.05, two-tailed student t-test). Statistically significant differentially expressed genes are denoted by red stars.

### 5. The marking of heterochromatic genes by H3K36me3 by dMES-4 is essential for heterochromatic gene expression

From our previous results, we were interested to know what other factors could be crucial for the regulation of heterochromatic gene expression. Earlier reports have suggested that the heterochromatic genes are marked by both H3K9me3 and H3K36me3 marks (Nicole C. Riddle et al. 2011). This is an interesting combination of functionally opposite histone modifications that have not been reported for other genes in the *Drosophila melanogaster* genome. Furthermore, the H3K36 methyl transferase MES-4 has been identified as an interactor of HP1a and co-localize to the pericentromeres (Alekseyenko et al. 2014), unlike the other known H3K36 methyl transferase dSet2. Therefore, we wanted to check the effect of dMES-4 downregulation on heterochromatic gene expression. In this regard, we analyzed the transcriptomic data for knock-down of dMES4 available in the public domain. We re-analyzed the data only for the heterochromatic genes both on the chromosome arms and unassembled heterochromatin (Het arms). We find that depletion of dMES4, results in expression of 23 heterochromatic genes being significantly down-regulated, **Figure 6**, including the well-studied heterochromatic genes like *light, Rolled, Nipped-A* (FDR adjusted p-value < 0.05) and more importantly, none of the heterochromatic genes was up-regulated. However, not all the down-regulated heterochromatic genes have > 1.5 fold change. Previous genetic studies have indicated that even 30% decrease in heterochromatic gene expression leads to discernable phenotypes (Lu et al. 2000) and thus we believe that these changes, though small, must be significant at the organismal level.

**Figure 6:**
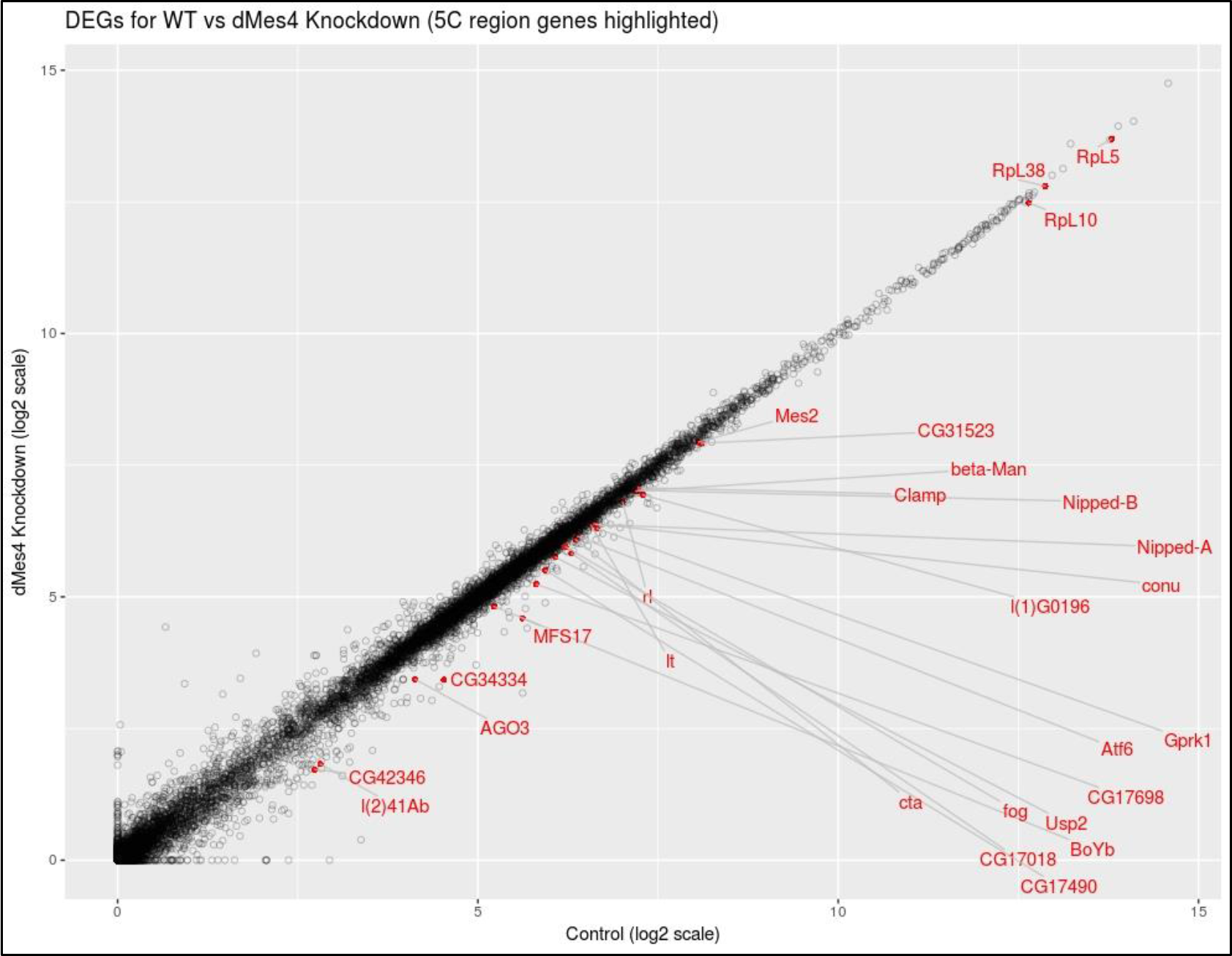
Knock-down of dMES4 down-regulates the expression of several heterochromatic genes-. Scatter plot showing the effect of dMES4 knock-down on the S2 cell transcriptome – comparison of WT vs dMES4 KD. Control RNA seq data on X-axis and dMES4 Knockdown RNA seq data on Y-axis. Significantly downregulated (present below the diagonal) heterochromatic genes, marked in red.

### 6. ADD1 facilitates expression of heterochromatic genes

HP1a and Su(var)3-9 maintains the heterochromatin structure and are also found to be binding at the heterochromatic genes (Greil et al. 2003; Nicole C. Riddle et al. 2011). However, given that these general heterochromatin factors bind all across the heterochromatin, the protein complex that assembles at the heterochromatic genes to mediate their transcription remains unknown. Furthermore, our results show that depletion of either HP1a or Su(var)3-9, the expression of heterochromatic genes is largely unaffected. This indicates that there is some factor that can interact with either HP1a or H3K9me3 marks to facilitate the expression of these genes. Thus, we chose to study dADD1 (CG8290) in this regard as it has been shown to interact with both H3K9me3 and HP1a and also colocalize at the pericentromeres ((Nicole C. Riddle et al. 2011). Therefore, it can probably mediate the regulation of heterochromatic gene expression in presence of either HP1a or Su(var)3-9 mediated H3K9me3 marks, thus, answering how the heterochromatic genes resists repression in spite the perturbations of genome organization upon depletion of either of these two proteins.

In order to understand how does dADD1 occupancy on the heterochromatic genes correlates with their expression, we grouped all the heterochromatic genes into 4 categories-a) No-expression-(0 FPKM) b) Low expression (1-10 FPKM) c) Medium expression (11-50 FPKM) d) High expression (>51 FPKM). We compared the enrichment of dADD1 (ChIP seq data) on each set of these genes along with the occupancy data of H3K9me3, H3K36me3 and HP1a. Interestingly, we find that for no and low expressing heterochromatic genes there is minimal enrichment of the dADD1 and H3K36me3 marks at the promoters and the gene body. On the contrary, there is increased binding of dADD1 upstream of the TSS, concomitant with an increased H3K36me3 binding at the gene bodies of the moderate and highly expressing heterochromatic genes, **Figure 7A**. This trend indicated that dADD1 could have a functional role in regulating heterochromatic gene expression as has also been previously suggested (Alekseyenko et al. 2014).

**Figure 7:**
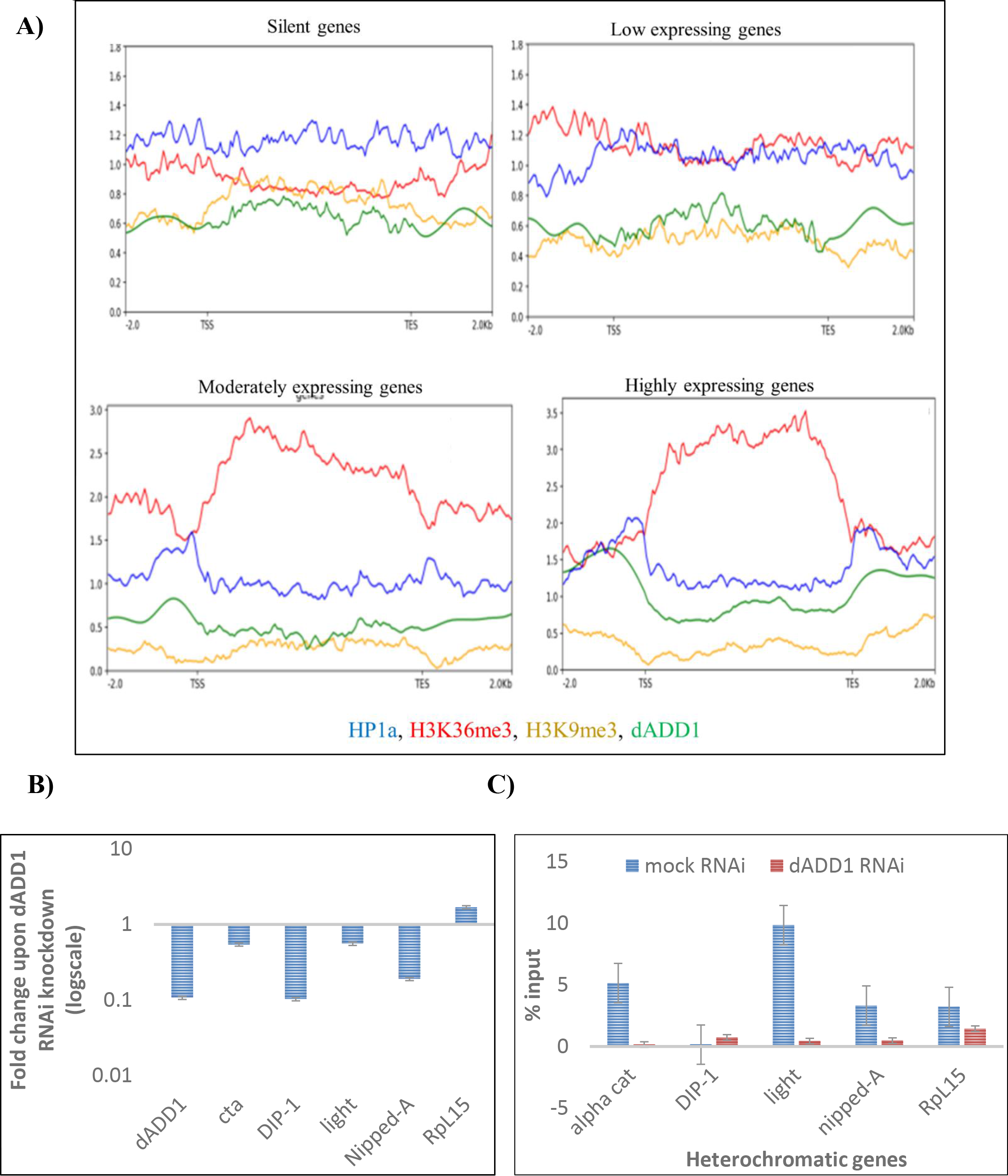
dADD1 is involved in regulating the deposition of H3K36me3 on the heterochromatic genes and their expression. **A)** Heterochromatic genes expression is associated with an increased enrichment of dADD1 at the TSS and H3K36me3 on the gene body **-** Correlation between expression levels of heterochromatic genes and occupancy of H3K9me3, H3K36me3, HP1a and ADD1 on the gene body across heterochromatic genes with no, low, moderate and high expression **B)** RNAi mediated dADD1 down-regulation results in lowering of expression of heterochromatic genes and H3K36me3 levels-*concertina, DIP-1, light* and *Nipped-A* show down-regulation and *RpL15* is not affected, (n=2). **C)** H3K36me3 levels (n=2) are also lowered on these heterochromatic genes upon dADD1 knock-down.

To confirm whether dADD1’s binding at the promoter of expressed heterochromatic genes has functional consequence on their expression, we performed RNAi mediated knock-down of dADD1 in S2 cells using dsRNA that targets the exons common to the three dADD1 isoforms. We confirmed the knockdown of ADD1 levels using qPCR. Next, we proceeded to check the levels of expression of heterochromatic genes. We checked the expression levels of candidate heterochromatic genes (Rp49 as an internal control), **Figure 7B**. We find that upon knock-down of dADD1 levels of 4 out of the 5 heterochromatic genes are significantly down-regulated. Next, we wanted to understand whether dADD1 knock-down affects the levels of H3K36me3 marks on the heterochromatic genes, **Figure 7C**. Hence, we performed chromatin immunoprecipitation using H3K36me3 antibody in dADD1 knock-down cells. The levels of H3K36me3 on the heterochromatic genes are lowered upon knock-down of dADD1. dADD1 has been shown to be involved in heterochromatin maintenance (López-Falcón et al. 2014; Chavez et al. 2017), however, our findings lead us to conclude that dADD1 is also an important player in regulating the expression of heterochromatin genes which is probably by facilitating the deposition of H3K36me3 at their exons.

## Discussion

### *Drosophila melanogaster* pericentromeres are organized into discrete TADs

One of the key findings of our study has been to map the long-range DNA interactions in the pericentromeric heterochromatin. We find that there exists a discrete TAD organization within the “HP1a/centromeric” domains that further compartmentalize local chromatin interactions. We observe a highly significant association of MARS with the Het TAD boundaries. Matrix associated Regions or MARs are the regions of the genome of that tethers and relocate a locus to the ribonucleoproteinaceous substratum of the nuclear matrix. Findings from our and other labs have shown that Nuclear matrix acts as the scaffold for various nuclear process (Kallappagoudar et al. 2010) and the organization of the genome through the MARs is crucial for modulating gene expression (Jenuwein et al. 1997; Wang et al. 2009). The overlap of the Het TAD borders with the S2 cell MARs is an experimental validation of their hypothesized role in pericentromeric genome organization (Saha, Sowpati, and Mishra 2018). Whether this means a functional correlation can be ascertained by further studies aimed at deleting the MARs in the genome and looking at the changes in genome organization and heterochromatic gene expression.

In lines with the reports for the global TAD organization (Ulianov et al. 2016) in *Drosophila*, we find that dCTCF and BEAF32 are enriched at the TAD borders. Intra-TAD interactions are marked by GAF and in flies it has been also shown to mediate gene activation (Chopra et al. 2008). ChrX shows a specific enrichment of CP190 and Su(Hw) at the TAD borders. We do not find the Het TADs to be specifically enriched in active marks as opposed to the intra-TAD, as has been reported for rest of the *Drosophila melanogaster* genome. Although there are speculations, that the repeat elements could be playing a role in genome architecture in the pericentromeric regions, there is no distinguishable difference in the enrichment of particular repeat class at TAD borders compared to the intra-TAD regions. Many of the 5C interactions were shown to map to the enhancers obtained from the STARR seq (Arnold et al. 2013) indicating they are most likely, the functional enhancer-promoter contacts. We also propose that the Het TADs marks distinct replication-timing domains within the generally late-replicating heterochromatin. Binding of HP1a has been shown to modulate replication timing in genomic-context dependent manner (Schwaiger et al. 2010) and the structural organization of the pericentromeres reflects the same.

Since the structural organization of the genome correlates to its functioning, we overlapped the RNA seq data of the S2 cells with the mapped TADs. We find that heterochromatic genes sharing the same TAD have similar expression levels. This indicates that they share the same regulatory network of long-range interactions and might be transcriptionally-coregulated. Interestingly, we also find that the majority of genes located at the TAD borders (19 genes) show high to medium level of expression, as has also been reported for the euchromatic regions. However, the heterochromatic genes within a particular TAD do not share any common gene ontology.

Pericentromere associated domains have so for been reported only in mice (Wijchers et al. 2015). Given the dearth of information regarding the hierarchical genome organization within the pericentromere, our findings provide a detailed characterization of the *Drosophila melanogaster* pericentromeric domains which sets the stage for probing into its functional role in heterochromatic gene expression.

### Global Het TADs are dispensable for heterochromatic gene expression

Classical genetics experiments have provided us with insights into the role of two heterochromatic factors-HP1a (Su(var)205) (Lu et al. 2000) and the H3K9me3 methyl transferase Su(var)3-9 (Yasuhara and Wakimoto 2008) in the regulation of heterochromatic gene expression. However, a detailed global understanding of their effects on these genes has not been reported. Upon knocking down these two proteins in the S2 cells, we find that the pericentromeric genome organization is perturbed marked by increased inter-TAD interactions in comparison to the intra-TAD interactions. This is more pronounced in the case of HP1a, which is the critical player in heterochromatin maintenance. This could also be because upon the knock-down of Su(var)3-9, other H3K9 methyltransferases (G9a and SETDB1) can compensate for its function. HP1a is not an architectural protein as the dCTCF or cohesin but our results indicate that it does contribute to both structural and functional maintenance of constitutive heterochromatin genome organization.

At the level of transcription, we find that HP1a RNAi and Su(var)3-9 RNAi affect 198 and 117 genes differentially across the genome and 101 genes are common among them. Notable examples of upregulated genes are *Trl (GAF)*, snRNA and snoRNA genes, *Rad50, HmgZ, Nup75, Su(var)2-10* and downregulated genes are *LamC* and heat shock proteins respectively, some of which are known interactors or targets of the two proteins. Interestingly only a small subset, 21 heterochromatic genes and 8 heterochromatic genes were differentially expressed in HP1a and Su(var)RNAi conditions respectively, which also includes certain non-coding transcripts like *CR41501, CR33294, 18SrRNA-Psi: CR41602*. Interesting some of the genes (example CG30440, Cht3, CG40006, CG40498) affected by HP1a/ Su(var)3-9 knockdown are the Group III genes, that is they have enrichment of only inactive marks in both the stages of highest and lowest expression (Saha, Sowpati, and Mishra 2018). This indicates that there exist different mechanisms of regulation of of different class of heterochromatic genes.

Previous genetic studies had used mutant alleles of *Su(var)205* (HP1a) to look for the effects of *Su(var)205* depletion on the *lt* and *rl* heterochromatic genes. They showed that there is a variegated silencing of light and rolled in *Su(var)205* mutant larvae evident by the reduction of *light-* dependent autofluorescent granules in Malpighian tubules. (Lu et al. 2000). We find that *lt* and *rl* are not downregulated significantly (FDR > 0.05) with respect to the entire transcriptome. It should also be noted that the difference in protein levels upon knock-down of HP1a leading to discernable phenotypes was only 1.5 fold in the genetic studies by Lu and colleagues. Such subtle changes are not picked up as significant in our genome-wide transcriptomic data but might be sufficient to result in phenotypes in the whole organism or on specific tissue type. However, our results are consistent with the report showing no change in expression of *light* and *rolled* upon RNAi mediated depletion of HP1a or Su(var)3-9 in Kc167 cells (Greil et al. 2003). Due to the drawback of the transient nature and <100% knock-down in RNAi conditions the fact that epigenetic landscape can be sustained for few cell-divisions to facilitate heterochromatic gene expression despite the pertubations in genome organization, cannot be ruled out. However, it is also a likely possibility that at least in the heterochromatin, the structural higher-order genome organization in the heterochromatin is a consequence rather than being the deterministic factor of heterochromatic gene expression. Several recent reports have suggested variable effects of disruption of genome organization on gene expression, for example, the drastic phenotypic effect of CTCF depletion to mild changes in gene expression upon acute CTCF depletion in cultured cells (Nora et al. 2017)(Kubo et al. 2017). On the similar lines, we believe that heterochromatic genes expression is regulated by more than one pathways. Consequently, the increase in inter-TAD interactions but not the intra TAD interactions indicates that the required local chromatin contacts are maintained for the intra TAD genes as opposed to the heterochromatic genes at the TAD borders being affected due to changes in the global heterochromatin structure in the HP1a RNAi condition. This is in line with the chromosomal translocation experiments where the severity of the *lt* gene’s variegation depended on where the chromosomal breaks were and how much of the dose of heterochromatic factors have been lost by the perturbation (Wakimoto and Hearn 1990). Thus, we posit that the local chromatin interactions within the TAD and heterochromatin dose provided by the heterochromatin factors are sufficient to sustain the expression of heterochromatic genes in the RNAi conditions. These findings provide a global perspective on the contribution of the pericentromeric hierarchical genome organization and heterochromatic factors (HP1a & Su(var)3-in modulating majority of the heterochromatic genes’ expression and validation of the candidate based genetic studies with modern genomics methods.

### dADD1 and dMES-4 in concert with heterochromatic factors regulate the permissiveness of heterochromatic gene expression from the pericentromeric heterochromatin

Riddle and colleagues had reported a comprehensive analysis of the heterochromatic epigenetic landscape in fly embryos, S2 and Kc167 cell lines. They find the heterochromatic regions harbour interesting combination of active and repressive epigenetic marks including the presence of H3K9me3, H3K36me3 and HP1a at the gene bodies of heterochromatic genes (Nicole C. Riddle et al. 2011). Interactome studies for HP1a proteins have shown that one of the interactors is H3K36me3 methyltransferase dMES-4, whose occupancy is enriched at the heterochromatic genes (Alekseyenko et al. 2014). This prompted us to look into the relationship of the transcription elongation mark H3K36me3 and the repressive marks of H3K9me3 and HP1a in the context of heterochromatic gene expression. We find that dMES-4 knock-down downregulates the expression of several heterochromatic genes most of which belong to Group-I (Saha, Sowpati, and Mishra 2018) or Group C/D as per (N.C. Riddle et al. 2011). These genes have the presence of H3K9me3, HP1a and H3K36me3 at their exons. These genes do not get affected upon single knock-down of either HP1a or Su(var)3-9 but are downregulated upon depletion of dMES-4. This indicates that the combinatorial histone modification of H3K9me3/HP1a and H3K36me3 on these genes is of functional relevance.

The epigenetic regulation of the heterochromatic genes is rather complex and our interest to learn about what other proteins could be regulating the epigenetic landscape of these genes led us to dADD1-*Drosophila* homolog of human ATRX. hATRX is an ATP-dependent chromatin protein in humans that belongs to the SNF2 family of chromatin remodeler consisting of an N-terminal ADD (ATRX-DNMT3-DNMT3L) domain. Mutations in the ADD domain causes the decreased DNA methylation, aberrant chromosome segregation and misregulated gene expression leading to mental retardation and thalassemia in humans (Iwase et al. 2011). The h*ATRX* gene diverged in *Drosophila* into dADD1 (ADD DNA binding domain but lacks the ATP helicase) and dXNP/ATRX (ATP helicase) (López-Falcón et al. 2014). In *Drosophila*, further biochemical characterization of dADD1 showed that all the three isoforms of it localize to the pericentromeric and telomeric heterochromatin and interacts with dXNP/ATRX (López-Falcón et al. 2014). *In vitro* assays have shown that dADD1 binds to H3K9me3/H3K4me0 and interacts with HP1a and acts as a mild suppressor of variegation in flies (Alekseyenko et al. 2014).

We find that there is an increase in the binding of dADD1 at the TSS and H3K36me3 at the gene bodies of the moderate to high expressing heterochromatic genes as compared to the silent or low expression. Our results show that dADD1 RNAi lowers the enrichment of H3K36me3 and also the expression of the heterochromatic genes. In the case of the human homologue hATRX, the association with the heterochromatin is established by localizing with H3K9me3/H3K4me0 and dADD1 recruitment at these genes is stabilized by HP1a–forming a tripartite interaction with HP1a and H3K9me3. dADD1 also recognizes this combinatorial histone mark. Thus when Su(var)3-9 is knocked down other H3K9 methyl-transferase can provide the required environment for dADD1 localization. While in case of HP1a knock-down condition, the stabilization of dADD1 binding at the heterochromatic genes is reduced due to the presence of the only H3K9me3 and thus more heterochromatic gene expression gets affected as compared to the Su(var)3-9 knock-down. We propose that dADD1’s ability to interact with both HP1a and H3K9me3 allows it to act as a sensor of the dosage of the heterochromatic environment on which the severity of compromised heterochromatic gene expression depends. We indirectly show that levels of H3K36me3 are irresponsible for the down-regulation of heterochromatic gene expression seen in case of dADD1 knock-down. Further experiments are required to dissect the inter-relationship of the role of dADD1, HP1a and dMES-4 in heterochromatic gene regulation. Even though, dADD1 and MES-4 were both identified as the interactor of HP1a and are enriched at the heterochromatic genes they were not found to interact with each other in the co-immunoprecipitation studies (Alekseyenko et al. 2014). Thus, there could be additional bridging factors that recruit dMES-4 at the exons when dADD1 and HP1a or H3K9me3 are present at the TSS, which is yet to be identified. While isolating the protein complex at the active heterochromatic genes remains challenging owing to the fact that some of these factors also binds to other regions of the heterochromatin, we believe that our findings have helped to identify the role of two new players in the regulation of the heterochromatic gene expression.

## Conclusions

Taken together, our results suggest that an interplay of the higher order chromatin organization and the epigenetic factors regulate the heterochromatic gene expression and that this process is regulated by more than one pathways. Het TAD structure compartmentalizes the similarly expressing heterochromatic genes where in the local chromatin contacts ensure their transcriptional regulation, **Figure 8.** dADD1 and dMES-4 are important *trans* factors that along with heterochromatic factors (HP1a and Su(var)3-9) maintain the epigenetic landscape that is distinct from the neighbouring heterochromatin and thus, is required for the expression of these genes. HP1a or Su(var)3-9 depletion affects the global heterochromatin architecture while local chromatin contacts and the combinatorial presence of one more heterochromatic factors (HP1a/H3K9me3) along with a gene activating factors like H3K36me3 is maintained.

**Figure 8:**
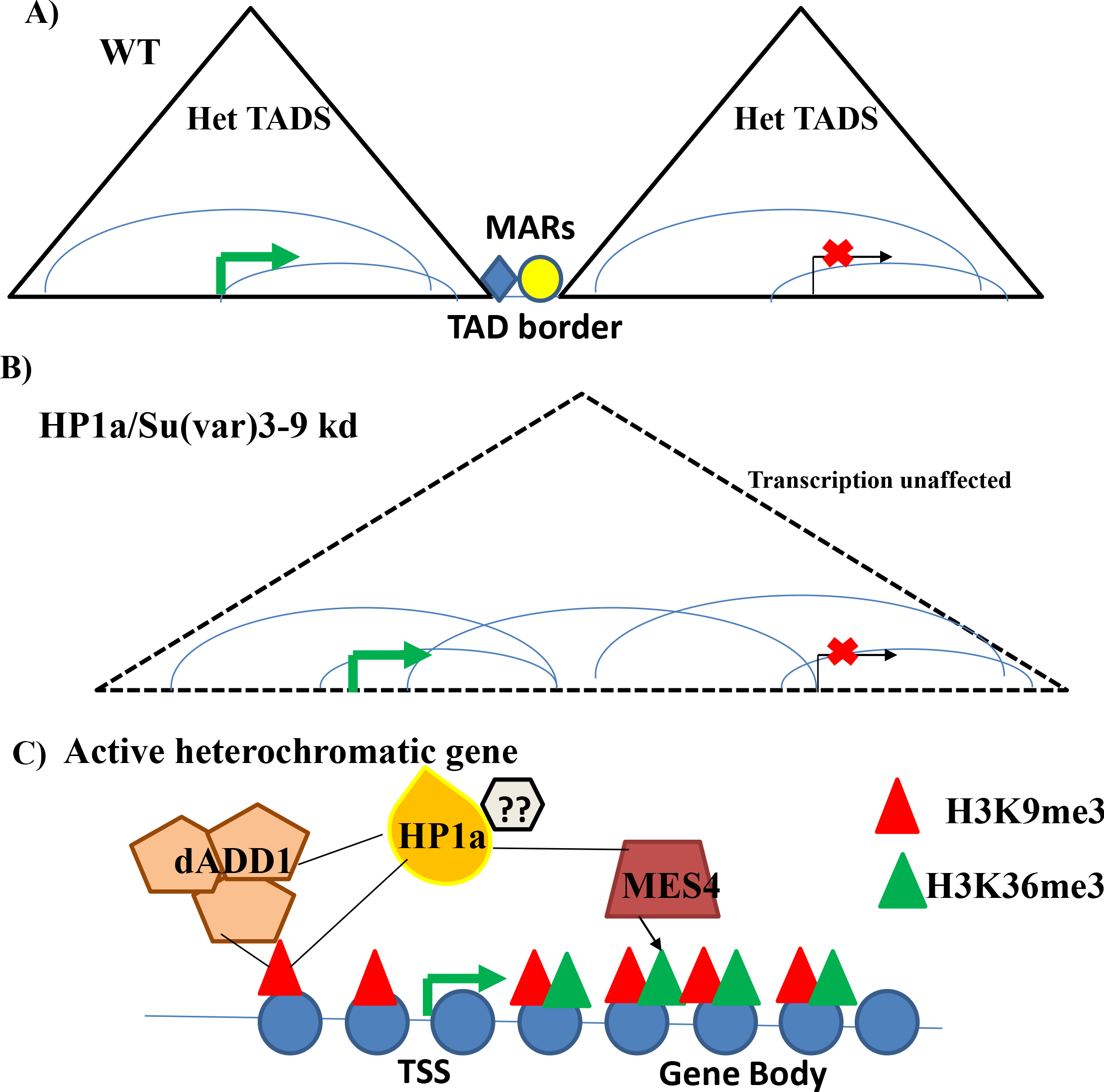
Interplay of pericentromeric genome organization and epigenetic landscape regulates heterochromatic gene expression-. **A)** Schematic representation of the pericentromeric heterochromatin divided into Het TADs that encompasses similarly expressing heterochromatic genes. **B)** Upon knockdown of heterochromatic proteins like HP1a or Su(var)3-9, the global TAD structure is perturbed but the local intra-TAD interactions maintained are sufficient to sustain heterochromatic gene expression. **C)** *Drosophila melanogaster* active heterochromatic genes show enrichment of dADD1 upstream of the TSS. dADD1 is probably involved in the regulation of heterochromatic gene expression in presence of heterochromatic factors (H3K9me3 marks and/or HP1a) and dMES4 is responsible for the deposition of H3K36me3 marks on the heterochromatic genes. The combinatorial histone mark of H3K9me3/HP1a and H3K36me3 at the exons facilitates the expression of heterochromatic genes marking them differentially from the surrounding repressive heterochromatin.

Further delineation of the molecular mechanisms in heterochromatic gene expression remains to be accomplished and so does the understanding of how has the evolution of Drosophilids shaped the epigenomic toolkit of heterochromatin to sustain the expression of heterochromatic genes in the repressive environment in only certain species. However, our findings provide some mechanistic insights into the role of important heterochromatin factors in regulating the variable chromatin landscape of the pericentromeres, thereby balancing the heterochromatic gene transcription and transposable element repression.

## Materials and methods

### Cell culture

S2 [*Drosophila* embryo (20-24hours)] derived cell-line was maintained in Schneider’s insect media with 10% heat-inactivated FBS and 1% Penicillin-Streptomycin at 25°C in the tissue culture incubator.

### 3C and 5C library preparation

#### Generation of 3C control library

The 3C and 5C libraries were prepared as described previously (Dostie and Dekker 2007). Purified genomic DNA from S2 cells was used for generating the 3C control library. 20μg of genomic DNA was digested at 37°C overnight using EcoRI (700U) restriction enzyme. Digested genomic DNA was purified using phenol: chloroform (1:1vol/vol). Digested DNA was resuspended and an aliquot for checking digestion efficiency. The ligation reaction was set up as follows: Digested DNA: 175μl; 10X T4 DNA ligase buffer-20μl, 10mg/ml BSA-2μl, 100mM ATP-2μl and T4 DNA ligase (300 cohesive ends unit per μl)-1μl and incubated at 16°C overnight. The digested and ligated DNA is the 3C control library and is suspended in 200ul TE. An aliquot was run on the agarose gel to verify digestion and ligation reactions.

#### Generation of the 3C library

5 × 10^7^ S2 cells were grown to confluency and 95-98% viability in the Schneider’s media for each of the 3 biological replicates. Cells were resuspended in 22.5ml of PBS/fresh media and proceeded for crosslinking using 37% formaldehyde strictly for 10 min. 1.25ml of 2.5M (5 min room temperature and 15 min on ice) glycine was used for quenching. Cross-linked cells were disrupted on ice using 15 strokes in a Dounce homogenizer (pestle A) without frothing. The cell lysate was centrifuged for 5min at 2,000g at RT and suspended in 500μl of 1X Restriction buffer 2 (NEB). This step was repeated thrice. The lysate was divided in 10 aliquots of 50 μl each. To each aliquot 337μl of 1X restriction buffer and 38μl of 1% SDS was added and incubated at 65°C for 10min exact (as prolonged incubation results in loss of cross-linked interactions). 44μl of TritonX-100 added to each tube to quench the SDS for facilitating ligation reaction. Finally, 400U of EcoRI (NEB) was added and incubated overnight at 37°C. 86μl of 10% SDS was added, mixed well and incubated at 65°C for 30 min. These 575μl reactions were used for ligation. Ligation reaction was set up as follows: Cell lysate-575μl; 10X T4 DNA ligase buffer (NEB)-745μl; 10% Triton X-100-745μl, 10mg/ml BSA-80μl, 100mM ATP-80μl; T4 DNA ligase (NEB): 10μl and water -5.96ml. Ligation was performed at 16°C for 2hours. Subsequently, 50μl of 10mg/ml Proteinase K was added and incubated overnight at 65°C followed by adding another 50μl of Proteinase K and another incubation with 2hours at 65°C the next day. To each of the ten ligation tubes, 8ml phenol was added, vortexed and centrifuged to collect the aqueous phase. The same procedure was repeated using 8ml of 1:1 vol/vol phenol: chloroform. To the aqueous phase, 3M sodium acetate pH 5.2 and chilled absolute ethanol was added to precipitate the 3C DNA. The library was dissolved in TE pH-8 and RNaseA treatment (1μl of 10mg/ml) was performed for 15min at 37°C. The 3C library should run as a tight band with molecular weight of more than 10kb, no or very little RNA and no undigested genomic DNA in the well. PCR titration was performed by using primers from a gene desert region in the *Drosophila melanogaster* genome (Cooper and Kennison 2011). Quality control of the 3C library was performed using positive interactions from reported interactions in the literature (H. B. Li et al. 2013). The 3C library generated was used for 5C library preparation or for PCR reactions to validate the 5C interactions. K1/K2 primers were used as internal controls(H.-B. Li et al. 2011).

### 5C Primer design and dilution

5C primers are designed using my5C tools for the pericentromeric regions: chr2L: 21900975-23011544, chr2R:1-1385689, chr3L: 22855576-24543557, chr3R: 1-428656, chrX: 21600796-22422827. Parameters used in primer design include: U-BLAST, 3; S-BLAST, 50; 15-MER: 800;MIN_FSIZE, 100; MAX_FSIZE, 50,000; OPT_TM, 65; OPT_PSIZE, 30. Primers were excluded wherever unique mapping was not possible with highly repetitive sequences. The universal T7 sequence was added to all forward primers (5’-TAATACGACTCACTATAGCC-3’) and the reverse complement to the universal T3 sequence to all reverse primers (5’-TATTAACCCTCACTAAAGGGA-3’). 5C forward and reverse primers are then pooled separately and diluted to the 50μM final concentration. The reverse primers are phosphorylated using Polynucleotide kinase (NEB).

### Conversion of 3C libraries to 5C libraries

On ice and in separate tubes, 3C and control libraries were mixed with salmon sperm testis DNA to a total DNA mass of 1.5μg such that the amount of 3C library taken represents approximately 150,000 genome copies to reflect the library complexity. 2.7μl of cold 5C primer mix containing 1μl of 10X 5C annealing buffer and 1.7fmol of each 5C primer and mixed well by pipetting. The samples were incubated at 95°C for 5min to denature the libraries and the primers. This was followed by incubating the mix at 48 °C for 16hours to anneal primers to libraries. 20μl 5C ligation buffer containing 10U Taq DNA ligase was added and incubated at 48°C for 1 hour to ligate 5C primers annealed to 3C junctions. The reaction is terminated by incubating at 65°C for 10min. No ligase, no template, and no primer controls are included in the PCR reactions set up to amplify the ligated 5C junction using universal T3 and T7 primers. PCR reaction was set up as follows: 5C ligation products: 6μl; 10X PCR buffer −2.5μl; 25nM dNTP mix-0.2μl, 20μM forward primer T7 and 20μM reverse primer T3-0.5μl each; 5U/μl platinum Taq polymerase-0.2μl and water:15μl. PCR cycle used was as follows: denature 95°C 5mins, Annealing-95°C, 30sec; 60°C, 30secs, and extension 72°C, 30sec for 35 cycles Final extension: 95°C, 30secs, 60°C, 30 secs and 72°C, eight mins. 5C libraries should appear at 100bp on the agarose gel with minimal smear and no bands in the controls. The libraries were run on E-gels (Thermo Fisher Scientific) and purified from there to allow a minimal loss in comparison to gel-elution, before proceeding for Illumina sequencing library preparation (as per the Illumina manual).

### RNAi mediated knockdown

For RNAi knock-down of HP1a, Su(var)3-9 and dADD1 proteins, primers were designed on the introns ranging in size from 300-1000bp. For proteins with isoforms, primers (T7 overhangs on both forward and reverse primers) were designed on the common introns to ensure knockdown of all the isoforms. Using the primers, the exonic regions are amplified and gel-eluted to use as a template for generating double-stranded RNA using T7 *in-vitro* transcription kit.

### *In vitro* transcription

1μg PCR template DNA was used as a template for the *in-vitro* transcription reaction as per the instructions of the Ambion MEGAscript kit. The RNA is extracted using phenol/chloroform and precipitated using isopropanol. An aliquot is used for checking the integrity and size of the RNA product on the agarose gel and quantification using NanoDrop spectrophotometer.

### S2 cell RNA transfection for RNAi mediated knockdown

S2 cell transfections were done using Qiagen Effectene Transfection Reagent. 10^6^cells/ml were plated in 6-well plates 24 hours before transfection. For each well 1μg of dsRNA prepared using *in-vitro* transcription kit was used. The cells were incubated at 25°C for 3 days for dsRNA mediated knock-down of the targeted protein. On the 4^th^ day, the knock-down was validated by Western blotting using a specific antibody against HP1a and Su(var)3-9 and using qPCR for dADD1 knock-down. The RNAi experiments were done in duplicates for 5C sequencing and triplicates for total RNA sequencing.

### RNA preparation for RNA seq libraries

S2 cells are harvested and pelleted. 1ml of TRIzol reagent is added per 10^6^ cells and pipetted to homogenize the cell-lysate. RNA precipitated using isopropanol, dissolved in DEPC treated TE buffer and checked for quality before proceeding for library preparation using Illumina RNA seq library preparation kit.

### cDNA preparation

The template RNA is treated with DNaseI (1:100) dilution to destroy any residual DNA during the RNA extraction procedure. The cDNA for reverse-transcriptase PCR is prepared using PrimeScript™ first strand cDNA Synthesis Kit from TaKaRa. The cDNA is appropriately diluted to proceed for analysis by quantitative PCR

### Chromatin Immunoprecipitation

1×10^6^ S2 cells are grown to confluence, and 95-98% viability is used for preparing the chromatin. The cells are scraped and washed twice with 1X PBS. The cells are then fixed using 1% formaldehyde for 10 minutes at room temperature. The formaldehyde is quenched using 2.5M glycine. The cells are washed again with 1X PBS thrice and proceeded for lysis. 100μl of 1X Lysis buffer containing 1% SDS, 1mM EDTA and 10mM Tris pH-8 and 1X protease inhibitors are added for each 10^6^ cell pellet. Lysis is allowed for 30 minutes on ice. Shearing of the chromatin in the range of 150-200bp is done using Pico Bioruptor in the Bioconductor tubes (2×10^6^ cells in 300μl) for 20 cycles - 30 seconds on, 30 seconds off. The chromatin is spun, and the supernatant is snap-frozen using liquid N2 for long-term storage at −80°C. An aliquot is saved for checking the sheared chromatin size (both on an agarose gel and by Bioanalyser). It is de-crosslinked by incubating the aliquot at 65°C for overnight followed by incubation at 56°C for 1 hour after adding 5μl of Proteinase-K. The DNA is precipitated using phenol/chloroform and checked using agarose gel electrophoresis. The chromatin immuno-precipitation was performed using Diagenode Low Cell ChIP kit. 5-10μg of chromatin and 1-3 μg antibody was used for each reaction. The ChIP-ed DNA was used for qPCR using SYBR green Master Mix on ABI quantitative PCR machine.

### 5C data analysis and peak calling

5C sequencing was done in triplicates and mapped to dm3/R5 genome build of *Drosophila melanogaster*. For all the libraries, because of common ends of all 5C primers, trimming of both 5' (1-20 bases) and 3' (80-100) ends was done with fastx_clipper from the fastx_toolkit. Reads of poor quality were removed using fastq quality_filter. Only reads with a minimum Phred score of 25 across 80% of the read was retained. Reads passing the quality filters were mapped to a custom reference genome made from all possible combinations of forward/reverse primers using bowtie2. Reads that mapped to multiple combinations were discarded. Interaction frequency lists were generated using a custom Perl script, and the frequencies were normalized to the total mapped read count. The normalization factors used are: EcoRI_control: 11.62, EcoRI_5C_rep1: 8.65, EcoRI_5C_rep2: 8.67. The normalized lists were further normalized to expected frequency using HiTC package in R. The resulting data was saved as pairwise interaction lists for each chromosome and sample. These pairwise interaction lists were uploaded to the my5C tool (http://my5c.umassmed.edu by Dekker lab) (Lajoie et al. 2009), and further analysis was done using my5C to generate the heatmaps.

### Domain calling using directionality index and HMM

The read normalized 10kb binned 5C pair-wise interaction files for each chromosome was saved from my5C web-tool. To define regions that contain TAD boundary, we implemented the Directionality Index method (Dixon et al. 2012; Crane et al. 2015) in our region of study. This method utilizes the fact that regions at the periphery of TAD are highly biased in their interaction frequencies and is based on chi-squared test statistic where the null hypothesis is that bins do not show biased upstream and downstream interactions. Directionality Index (DI) calculation was performed using an R script, in a 100 kb X 100 kb square along the diagonal of the interaction frequency matrix for each replicate using the following formula:

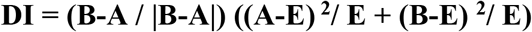

Where:

**A** is the sum of all interactions from a given 10 kb bin to the upstream till 100kb
**B** is the sum of all interactions from a given 10 kb bin to the downstream till 100kb
**E** is the expected number of interactions for each bin under the null hypothesis and it equals (A+B)/2

To predict TADs after estimation of DI, a Hidden Markov Model was constructed. The previously document script available in MATLAB (Crane et al. 2015) was rewritten in R language using CRAN package HMM. TADs were predicted across the replicates for each chromosome except for chr3R (because of too few interactions mapped, the HMM did not pick up any signals). The TAD boundary called for each replicate was pooled for a consensus boundary definition. For this purpose, a combined distribution plot for the distance between boundary calls for all chromosomes was plotted. Boundaries within 50kb of each other among replicates were considered consistent (>72.3% of all replicate boundary calls overlapped within this window). Using these parameters, TADs and TAD borders were defined for each chromosome.

### Defining inter and intra TAD interaction for comparing interaction in knockdown conditions

The read normalised pairwise-interaction matrices were taken from my5C web-tool. A grouped bar plot showing inter-TAD and intra-TAD interaction frequency were plotted. For this purpose, the genomic loci defined as TAD from our analysis, for each chromosome, were taken into account to calculate inter-TAD and intra-TAD interaction. The replicates (n=2 for knock-down samples and n=3 for Control) were pooled while making final inter- and intra- TAD interaction tables by taking a mean value of interaction score.

### Finding overlaps – intra-chromosomal and inter-chromosomal interaction datasets

The BEDTools suite (Quinlan and Hall 2010) was used to compute overlap of various genomic and epigenomic features with the intra-TAD and TAD boundaries. The BED file for each chromosome (intra-chromosomal interactions) was binned into 10kb windows (5kb for chrX). The number/frequency of overlap of each genomic feature with each 10kb bin was calculated. To check for any biases creeping up in our analysis, we went 500kb upstream/downstream on chromosome arms L and R respectively. The results were plotted as a heat map for each chromosome with features on the y-axis and 10kb bins of genomic co-ordinates on x-axis using the ggplot2 package in R. The TAD borders defined previously were also highlighted in the heat maps for intra-chromosomal overlaps. To calculate the statistical significance of our computed overlaps, regionR (DOI: 10.18129/B9.bioc.regioneR) Overlap Perm package was used for Permutation test, calculating the p-value for the numbers of overlaps of a particular feature with TAD borders as opposed to the randomized dm3 genomic regions were calculated and plotted in a graph. Various feature files were obtained from modENCODE/respective publications (Roy et al. 2010) and are listed in Supplementary Table

Similarly, for trans or inter-chromosomal interactions, a BED file was prepared to include all the unique anchor points, 10kb binned, across the two replicates to represent inter-chromosomal interaction. This was then intersected with the various genomic features, repeats, boundary elements and histone modifications as done for intra-chromosomal interactions. The overlaps of features in different combinations were visualized as an UpSet plot (Conway, Lex, and Gehlenborg 2017).

### Transcriptome analysis

Gene expression was quantified using paired-end (read length-151bp) RNA-Seq for Control, HP1 and Su(var) knockdown samples in triplicates. A preliminary quality check on data for finding errors in library preparation or sequencing was done using FastQC (version 0.11.5). The adapter removal was done using Cutadapt (version 1.11) (Martin 2011) in a paired-end mode with Phred score cut-off as 30 for both 3’ and 5’ ends and minimum read length to be kept as 50 bases. 82-90% reads passed this quality cutoff. The reads were then aligned to the reference genome, i.e., *Drosophila melanogaster* (build dm3) using STAR (Spliced Transcripts Alignment to a Reference)-Aligner (version 2.4.5a) using two-pass mode (Dobin et al. 2013). The Pearson Correlation Coefficient (PCC) showed that replicates correlate well among themselves for each condition (r>0.96 for all, the cutoff is >0.90). The quantification of gene expression into FPKM (Fragments per kilobase per million) and TPM (Transcripts per million) followed by differential expression was done using RSEM (RNA-Seq by Expectation Maximization version 1.3.0) (B. Li and Dewey 2011).

To see if genes falling in the same TAD follow a similar expression profile, we took the quantification step output files of RSEM method. From it, FPKM values across replicates for each condition was made into a single table and genes falling within our experimental region were a subset for all further analysis. A mean for FPKM value (combining the individual replicate’s FPKM value) was calculated for each gene and categorized into gene expression subgroups as followed in FlyBase. However, we categorized the Low, Medium and High expression categories mentioned are as follows: a) No-expression-0 FPKM b) Low expression 1-10 FPKM c) Medium expression 11-50 d) High expression >51. This criterion was also used for comparing gene expression with dADD1, H3k36me3, H3K9me3 and HP1a ChIP data. The differential gene expression analysis for KD samples against a control sample with FDR (False Discovery Rate) cutoff of 0.05 was used to retrieve differentially expressed genes.

### dMES4 knockdown analysis

The RNA-Seq for knockdown of histone methyl transferase dMes4 and its appropriate control was taken (Lhoumaud et al. 2014). The alignment using STAR-aligner, finding differentially expressed genes (DEGs) with FDR cut-off (0.05) using RSEM was done as mentioned above. The DEGs falling in our region of the study were shortlisted from this set.

### Replication timing data

The replication timing dataset for *Drosophila melanogaster* (build dm3) was taken from http://www.replicationdomain.com (Weddington et al. 2008). This was overlapped and plotted as an area plot over the identified TADs for our study with replication timing on the y-axis and genomic co-ordinates on the x-axis and visualized graphically.

### Additional files

**Additional File 1 (.xls):** Sequencing statistics of 5C and RNA seq datasets

**Additional File 2 (.xls):** Differentially expressed genes in HP1a KD and Su(var)3-9 KD

**Additional File 2 (.xls):** Differentially expressed genes in dMES-4 KD

## Supplementary Information (.pdf)

**NGS datasets of 5C-seq and RNA seq experiments:** GSE XXXXXXX (NCBI_SRA database)

## Acknowledgements

The authors thank RKM lab members for discussions and critical reading of the manuscript. PS acknowledges University Grants Commission (UGC), India for the doctoral fellowship. This work was supported by grants from Council of Scientific and Industrial Research (CSIR), India to RKM.

