## Supplementary material for "Interplay of pericentromeric genome organization and chromatin landscape regulates the expression of *Drosophila melanogaster* heterochromatic genes"

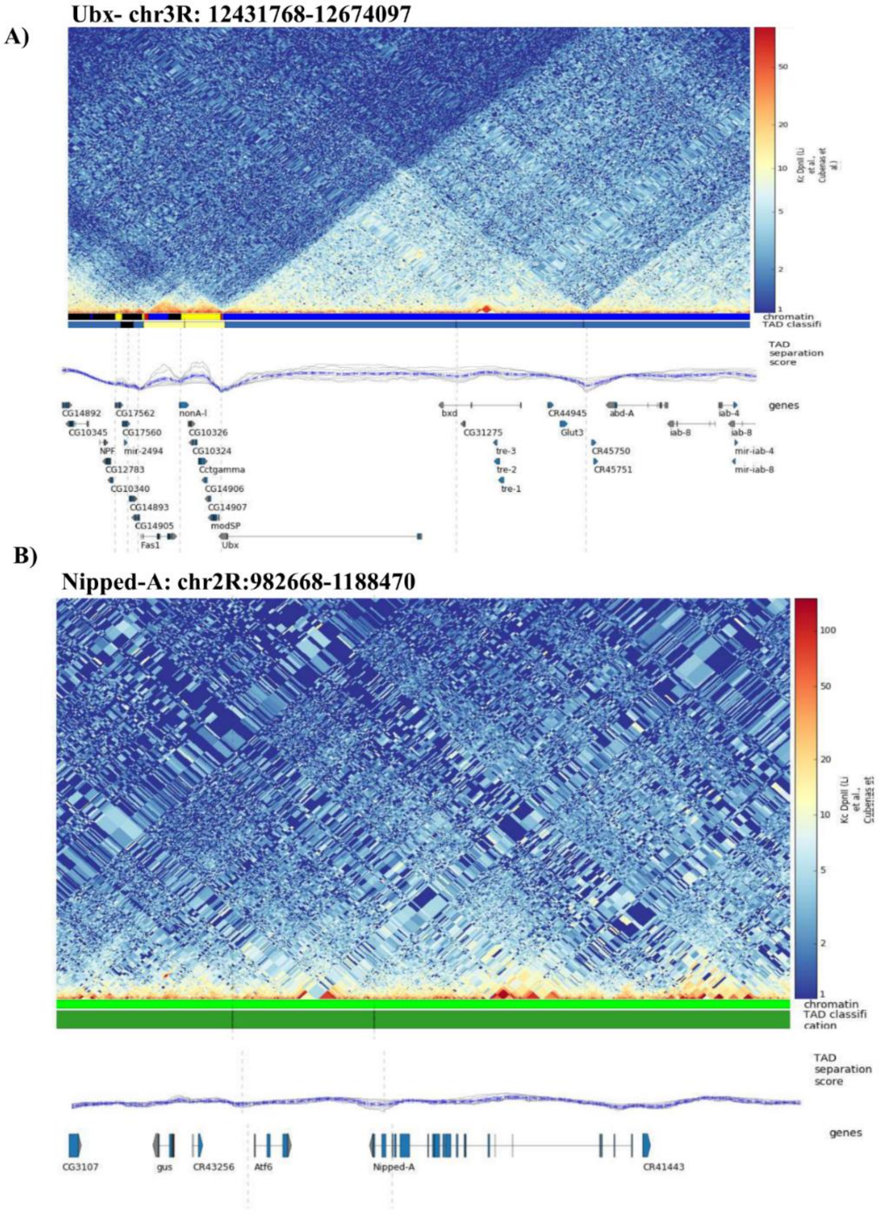


**Supplementary Figure S1: Heterochromatic regions lack sufficient information about the pericentromeric genome organization- A)** Snapshot of a euchromatic region of *Ubx* gene, The first panel depicts the TAD structure reported in Kc cells using Hi-C (Li et al &Cubentas Pott et al), followed by the TAD separation depicted in blue dotted lines shows dips at the TAD borders. The last track is for the genes. Ubx locus has well defined TAD information and TAD borders are indicated by the dips in the TAD score track. **B)** Snapshot for a heterochromatic gene *Nipped A* has limited TAD information The figures are generated using Chorogenome Navigator (http://chorogenome.ie-freiburg.mpg.de).

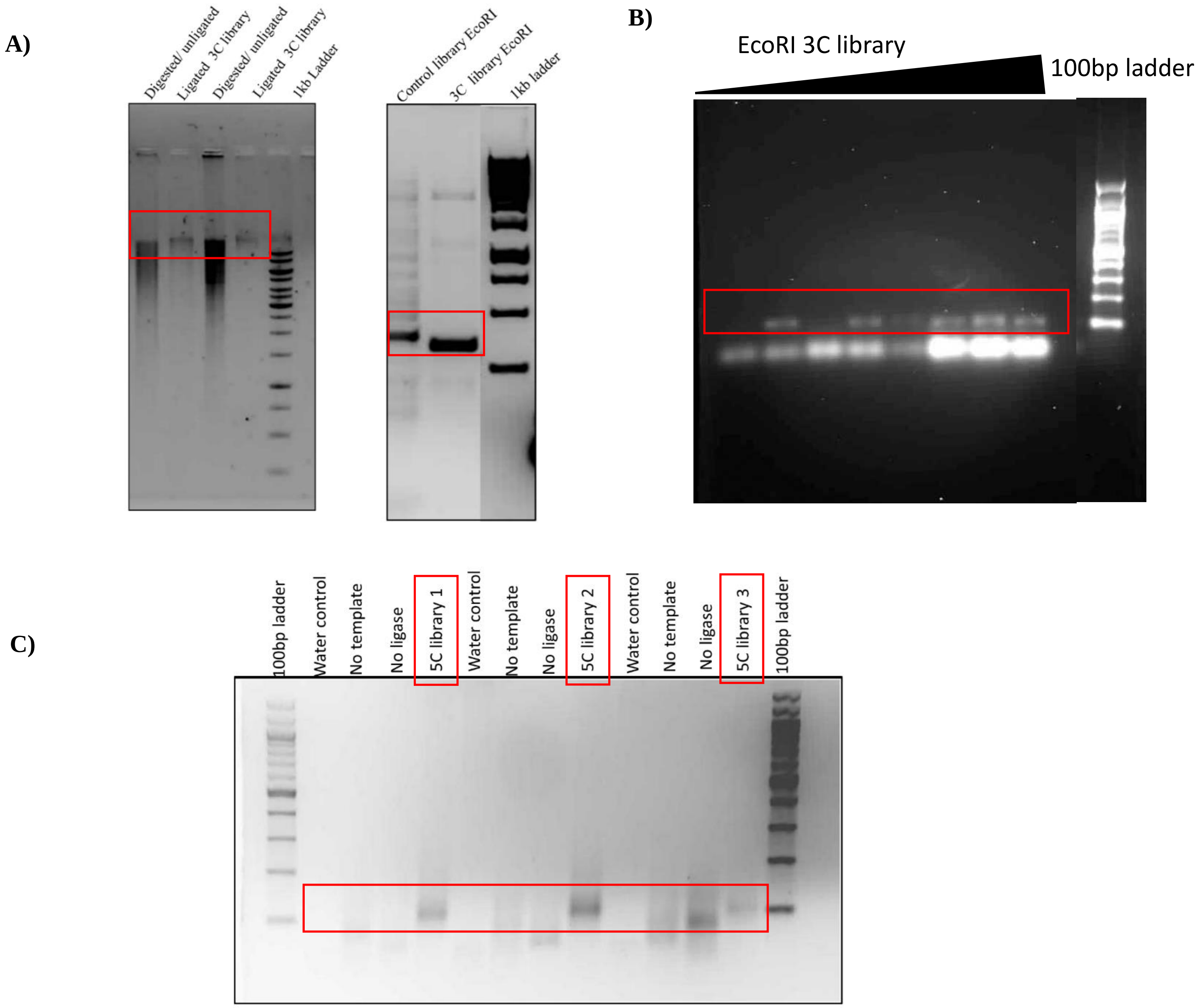


**Supplementary Figure S2: Quality control for 3C and 5C libraries** **A)** The agarose gel image on the left, shows the digested chromatin which is a smear but retracts to a tight band that runs above 10kb upon successful ligation. The gel image on the right shows PCR with positive control primers selected from published literature; the control 3C library gives multiple bands as opposed to the single prominent band for the 3C library. This indicates that the 3C library preparation is good and has the representation of the already reported interactions. **B)** Gel image showing titration of 3C library using primers designed at the gene desert region of *Drosophila* genome. **C)** The agarose gel image of the 5C PCR reaction where no template, no ligase, and no primer controls are taken for the replicates to ensure that the 5C library amplified is free of spurious PCR amplifications. The 5C library is 100bp in size as the primers are 50bp each (30bp complementary to sequence adjacent to the EcoRI site from the 5C region of interest including half of the EcoRI site and 20bp for the overhangs. The PCR products must have intact EcoRI site in the middle).

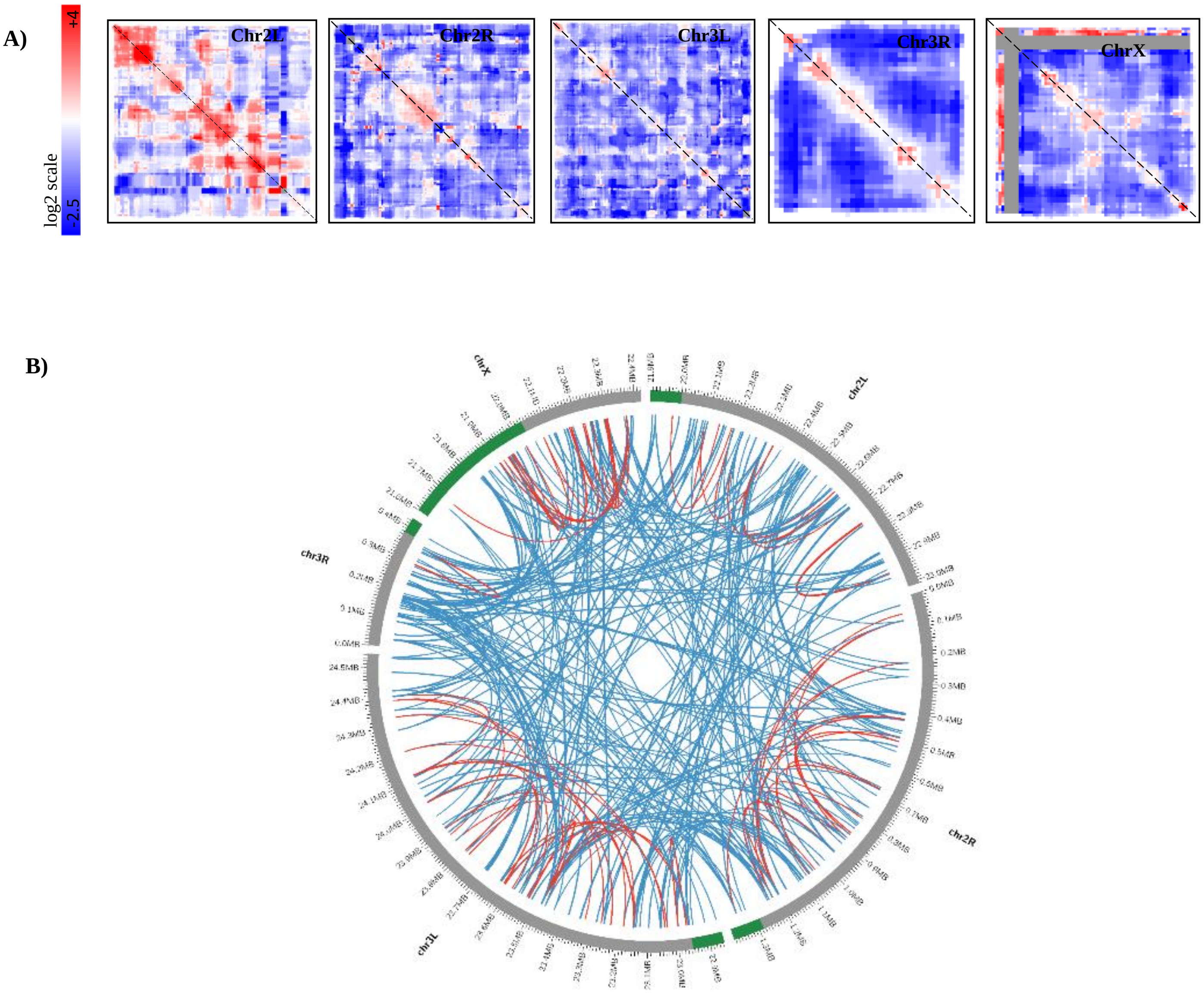


**Supplementary Figure S3: A)** Heatmaps for the 5C interactions mapped across 5 chromosomal arms- Chr2L, Chr2R, Chr3L, Chr3R and ChrX. The data is binned at 10kb (except Chr X at 5kb) and for each bin, the Interaction Frequency (in log2 scale) is plotted. The white bar in the heatmap of Chr X is indicative of the region where no sequence information was available and no primers could be designed. For all arms, euchromatic regions included in our study, partitions into a centromere-distal domain (E) distinct from the centromere-proximal domains of heterochromatin **C)** Circos plot of the inter-chromosomal interactions (blue) between the pericentromeric regions (green) of the chromosomes. Intra chromosomal interactions in red.

A)

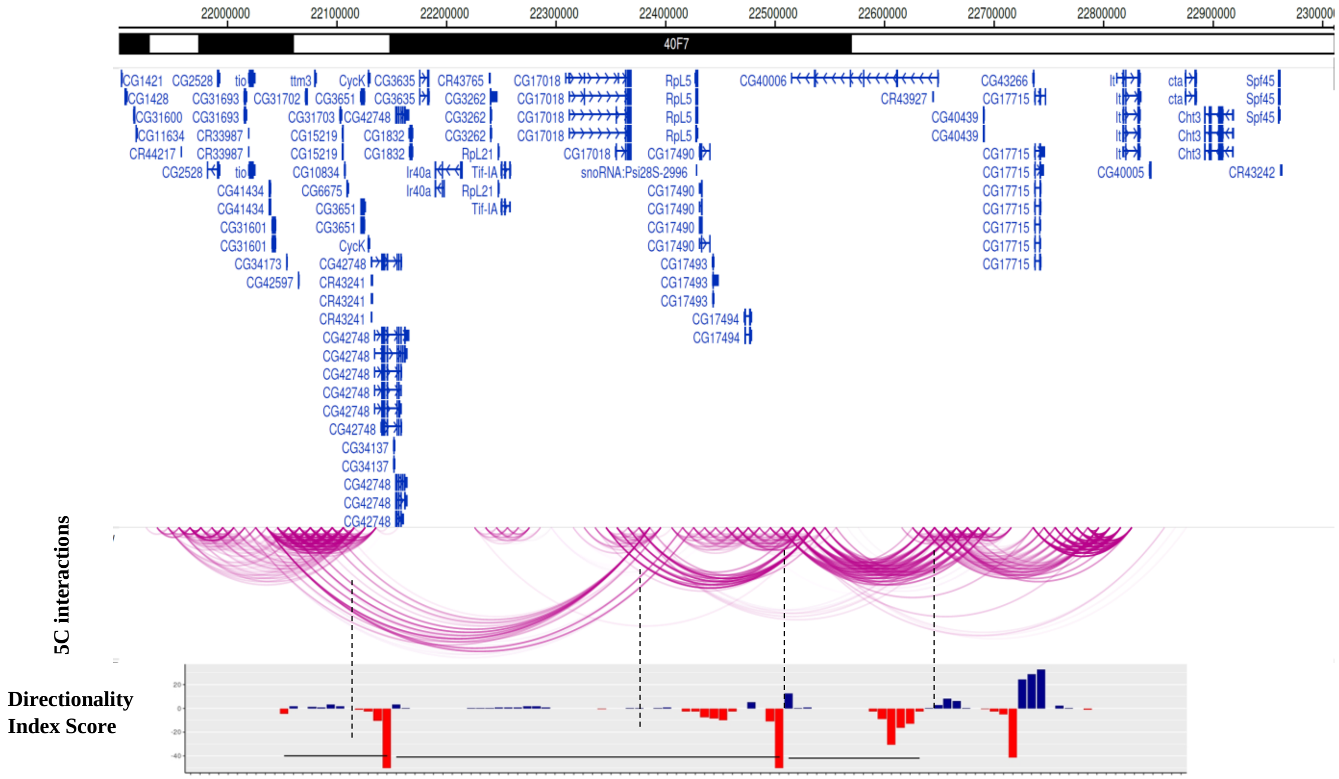


B)

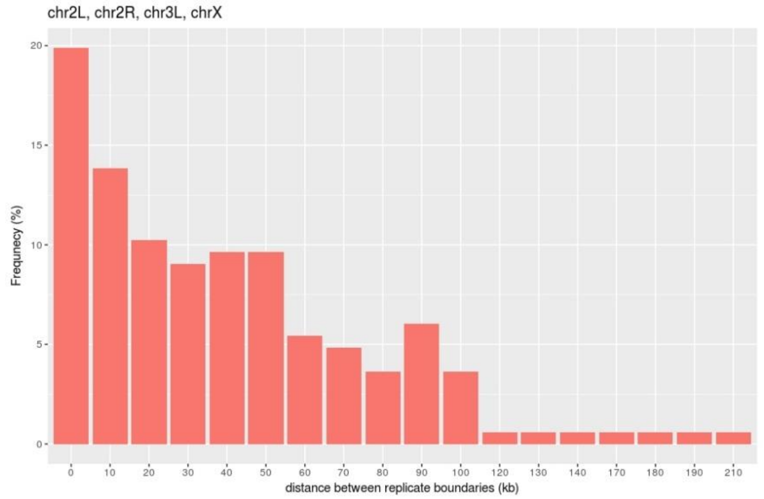


**Supplementary Figure S4: TAD calling by Directionality Index-** Epigenome Browser snapshot of Chr 2L:21900975-23011544 where the polytene bands are shown in black, genes in blue, long-range interactions mapped by 5C sequencing are represented as pink arc diagrams and the score of Directionality Index in bar plot the y-axis of directionality index score is not visible. Borders of self-interacting regions (dashed-line) show a drop in DI score indicating the directionality of the interactions. These scores are further used to determine the TAD borders computationally by Hidden Markov Model. **B)** Combined distribution plot for the distance between boundary called for all chromosomes. 50kb was defined as the final boundary zone for pooling of data as >72.3% of replicate boundary calls overlapped within this window. Using this as a window, TADs and TAD borders were defined.

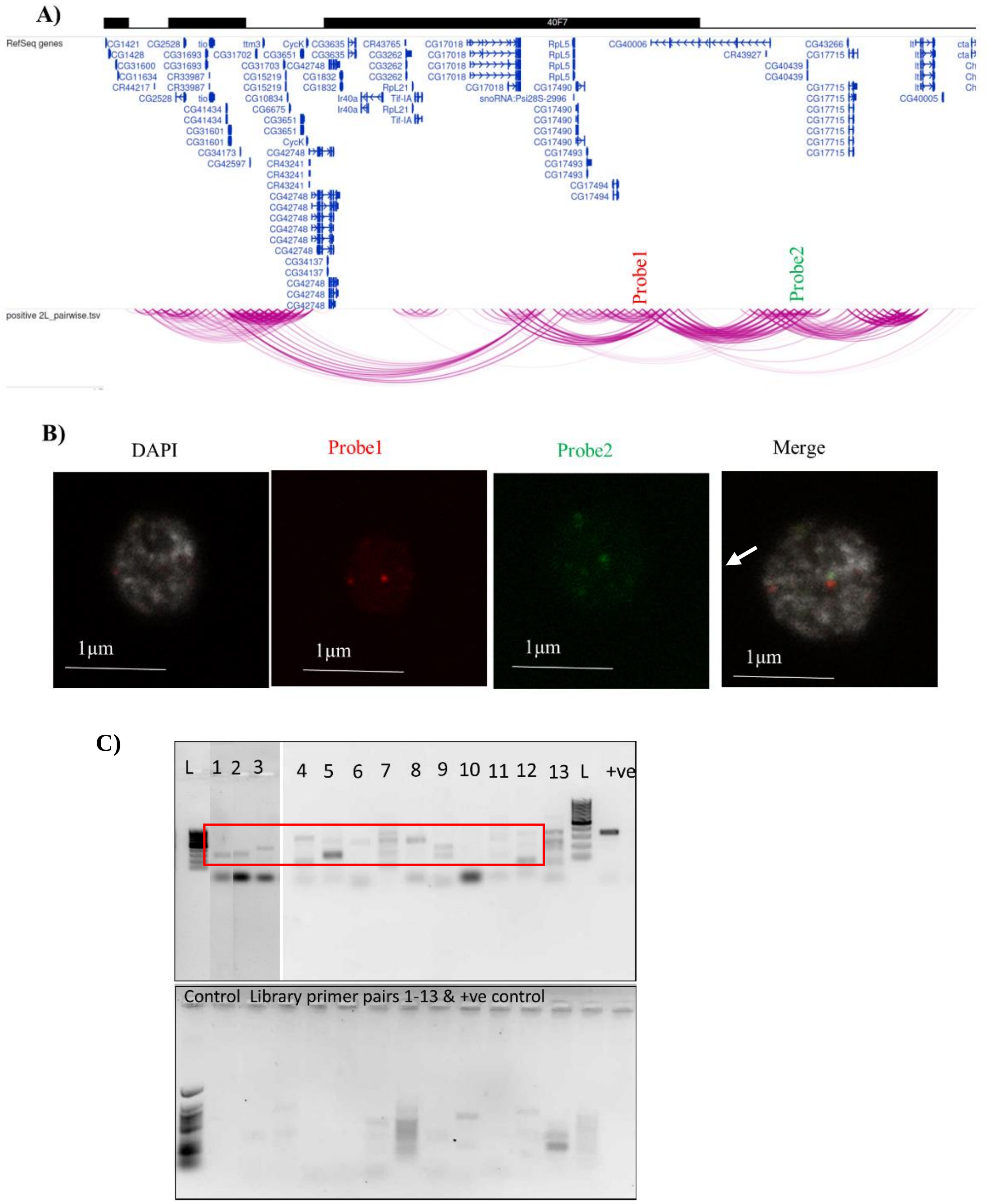


**Supplementary Figure S5: Validation of the 5C data- A)** FISH probes designed at the two ends of the TAD at Probe1 at Chr2L: 22346057-22356057 tagged by Alexa Fluor 488 and Probe 2 at Chr2L: 22636057-22646057 by Alexa Fluor 555, which are 280kb apart on the linear genomic distance. **B)** FISH shows that the two probes designed at two ends of the TAD come together in the 3D nuclear space (n=2) as evident from the merging of the red and green FISH signals. The nucleus is stained by DAPI and artificially colour changed to grey. **C)** Validations of 5C data using 3C PCR against 13 interacting regions from 5 chromosome arms label the gel lanes. The amplification of the product is pronounced in the 3C library and not in the control library.

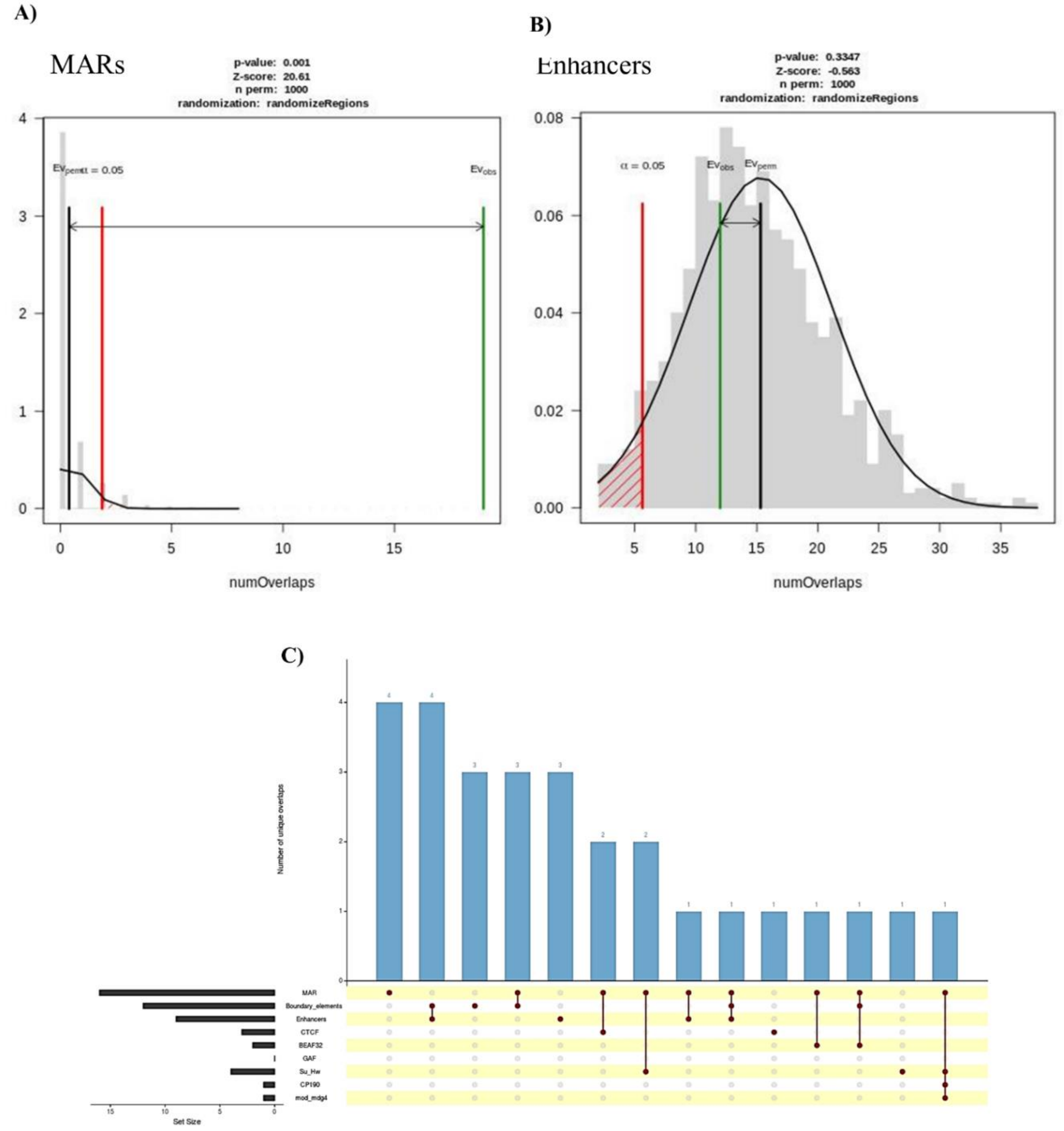


**Supplementary Figure S6: A)** Overlap permutation test to check the statistical significance of TAD borders overlapping with the MARs (***p value 0.001 is it = or <) and **B)** with the STARR seq enhancers (p = 0.3347) **C)** UpSet plot showing the distribution of overlap of Het TAD borders with various features.

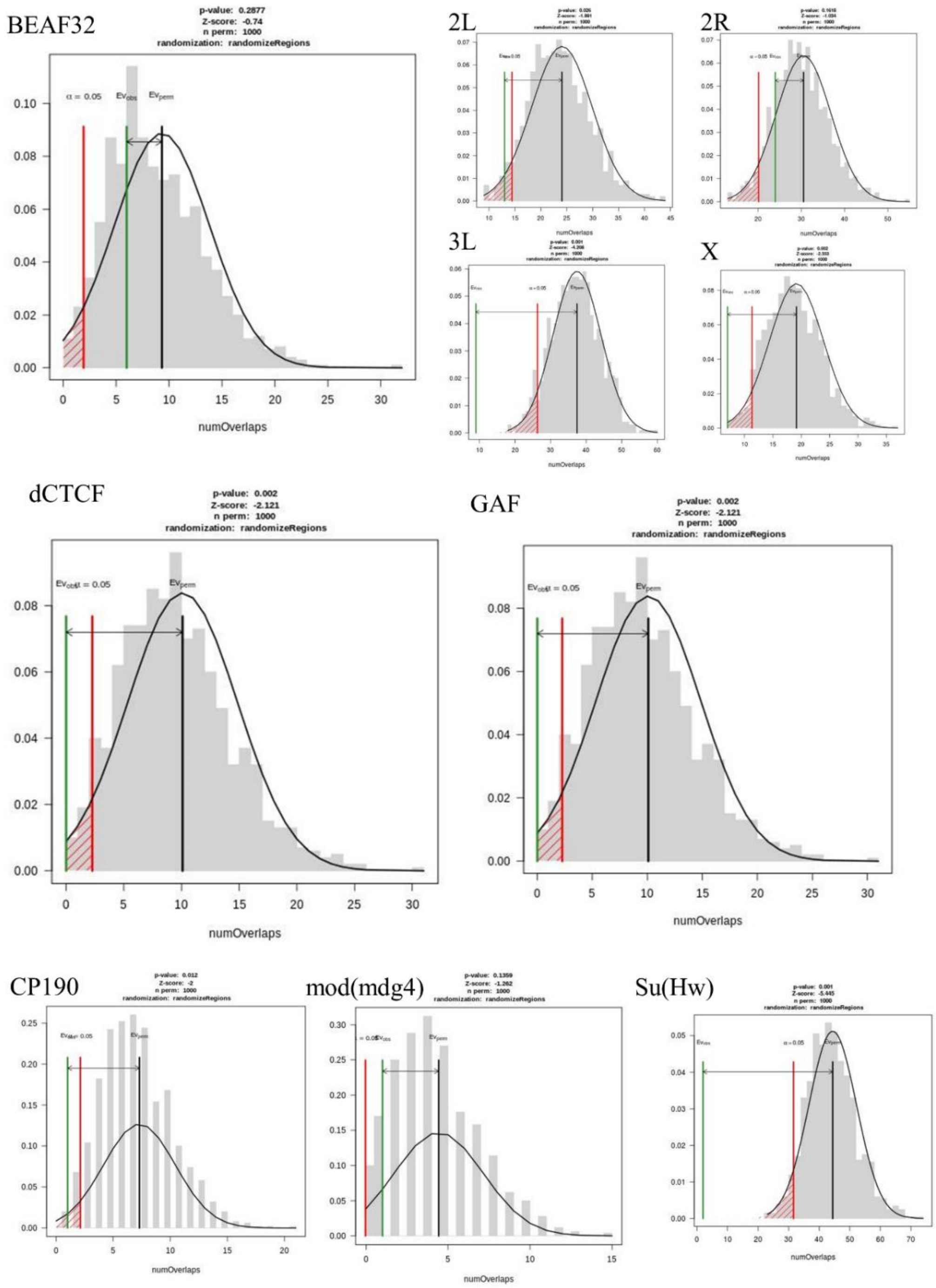


**Supplementary Figure S7: Overlap permutation statistical test for overlap of Het TAD borders with enrichment for binding of various architectural/ insulator proteins** -BEAF32 is most enriched at the Het TAD borders (combined p-value = 0.2, p-value for overlap across chromosome arms). dCTCF is the next most abundant protein binding at the Het TAD borders (See Supplementary Fig S5C). dTCF and GAF have the same pattern of enrichment (Fig 2). CP190, mod(mdg)4 and Su(Hw) (Supplementary Fig S5C) mark few TAD borders (only at Chr X).

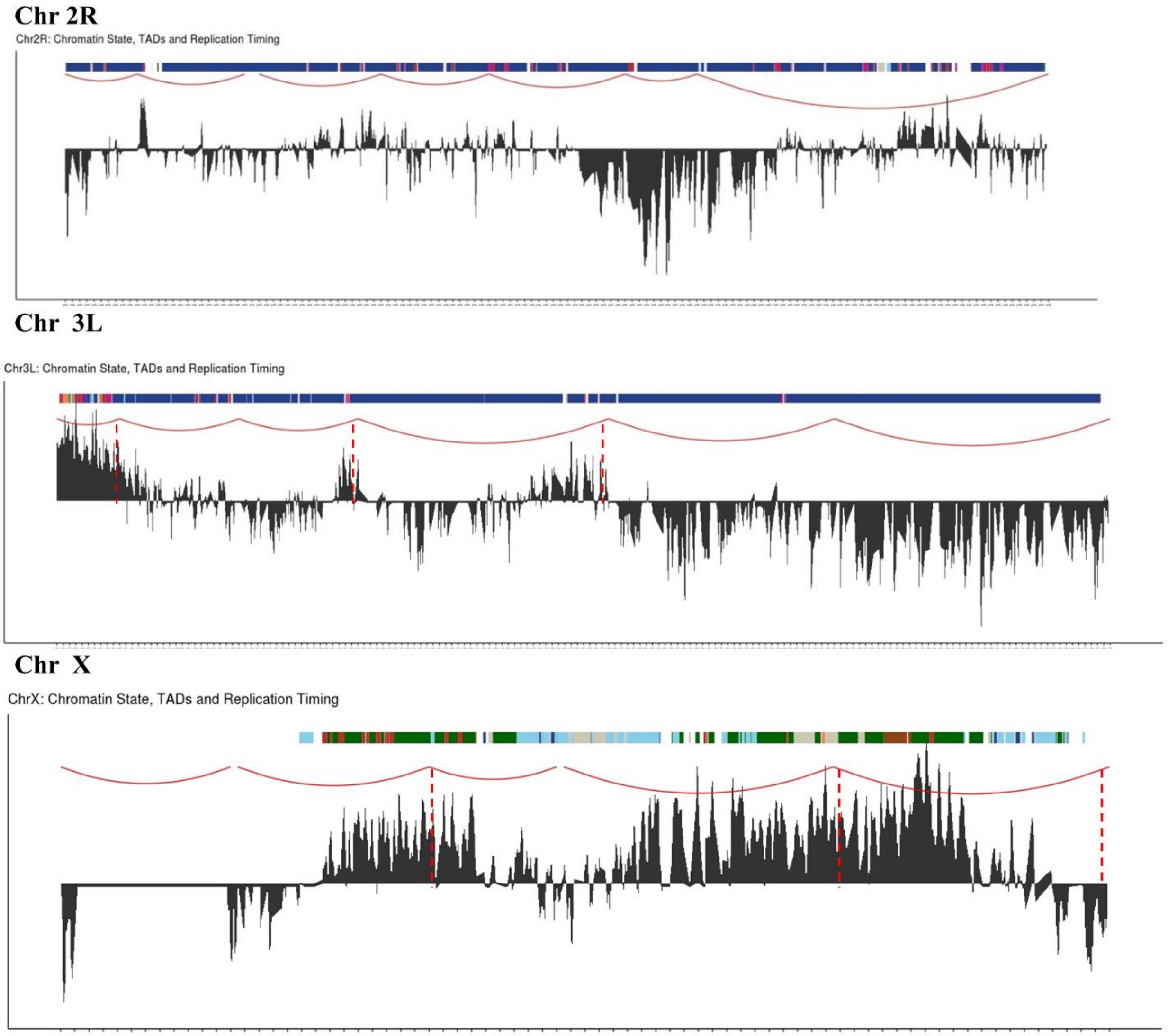


**Supplementary Figure S8**: Het TADs also demarcate replication timing domains in S2 cells. Overlay of the replication timing data (black peaks) with long-range interactions in red. Colour panel above denotes the 9 state chromatin model of *Drosophila melanogaster*

**Table 1**: List of TAD and TAD boundary coordinates

| **TADs** | | | **TAD boundaries** | | |
|---|---|---|---|---|---|
| TADs | | | | | |
| chr2L | 21900975 | 21996057 | chr2L | 21996057 | 22016057 |
| chr2L | 22026057 | 22116057 | chr2L | 22116057 | 22136057 |
| chr2L | 22136057 | 22536057 | chr2L | 22536057 | 22556057 |
| chr2L | 22556057 | 22686057 | chr2L | 22686057 | 22706057 |
| chr2L | 22706057 | 23011544 | | | |
| | | | chr2R | 109746 | 129746 |
| chr2R | 1 | 109746 | chr2R | 259746 | 299746 |
| chr2R | 129746 | 259746 | chr2R | 449746 | 469746 |
| chr2R | 299746 | 449746 | chr2R | 599746 | 619746 |
| chr2R | 469746 | 599746 | chr2R | 789746 | 809746 |
| chr2R | 619746 | 799746 | chr2R | 889746 | 909776 |
| chr2R | 799746 | 899746 | | | |
| chr2R | 909776 | 1385689 | chr3L | 22956370 | 22976370 |
| | | | chr3L | 23146370 | 23166370 |
| chr3L | 22855576 | 22956370 | chr3L | 23336370 | 23356370 |
| chr3L | 22976370 | 23146370 | chr3L | 23736370 | 23756370 |
| chr3L | 23166370 | 23336370 | chr3L | 24006370 | 24116370 |
| chr3L | 23356370 | 23736370 | | | |
| chr3L | 23756370 | 24006370 | chrX | 21779812 | 21789812 |
| chr3L | 24116370 | 24543557 | chrX | 21914812 | 21924812 |
| | | | chrX | 22004812 | 22019812 |
| chrX | 21600796 | 21779812 | chrX | 22199812 | 22209812 |
| chrX | 21784812 | 21919812 | | | |
| chrX | 21924812 | 22004812 | | | |
| chrX | 22019812 | 22199812 | | | |
| chrX | 22209812 | 22422827 | | | |

| Genes falling in TAD borders (low expression genes in grey) |
|---|
| 2L: CG31693, CG3651, CG15218, CG40006, CG40439 |
| 2R: CG17665, CG40129, CG40293, CG41440, CG17704, CG33492 |
| 3L:CG32230, CG17698, CG40045, CG40452 |
| X:CG17600 CG32499 |

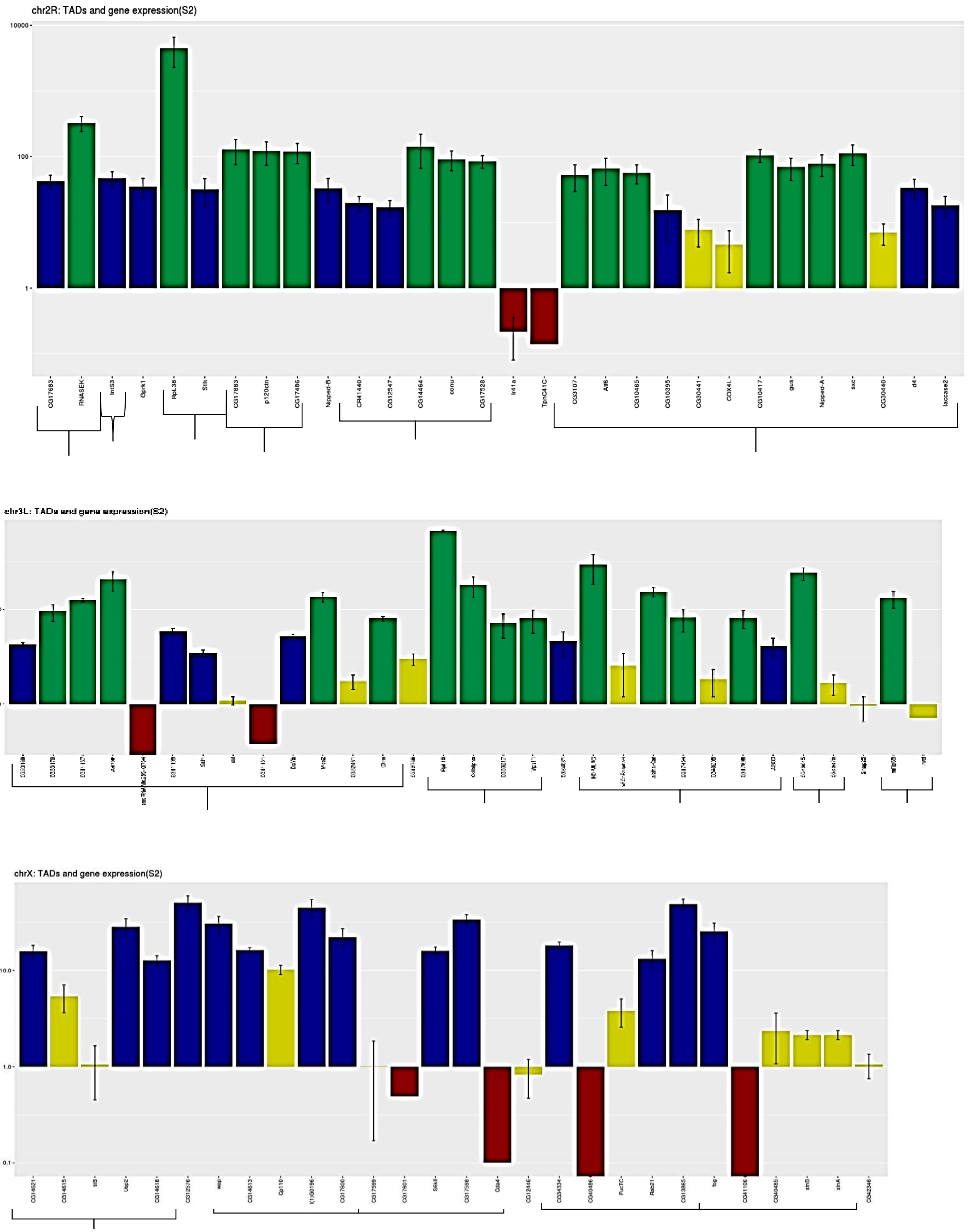


**Supplementary Figure S9: Expression pattern of heterochromatic genes in S2 cells correlates with the het TAD organization**: Bar plots showing the transcriptomic data for the S2 cells (3 biological replicates- Mean +SE) and their correlation with the Het TAD organization indicates the role of pericentromeric genome organization in their transcriptional regulation.

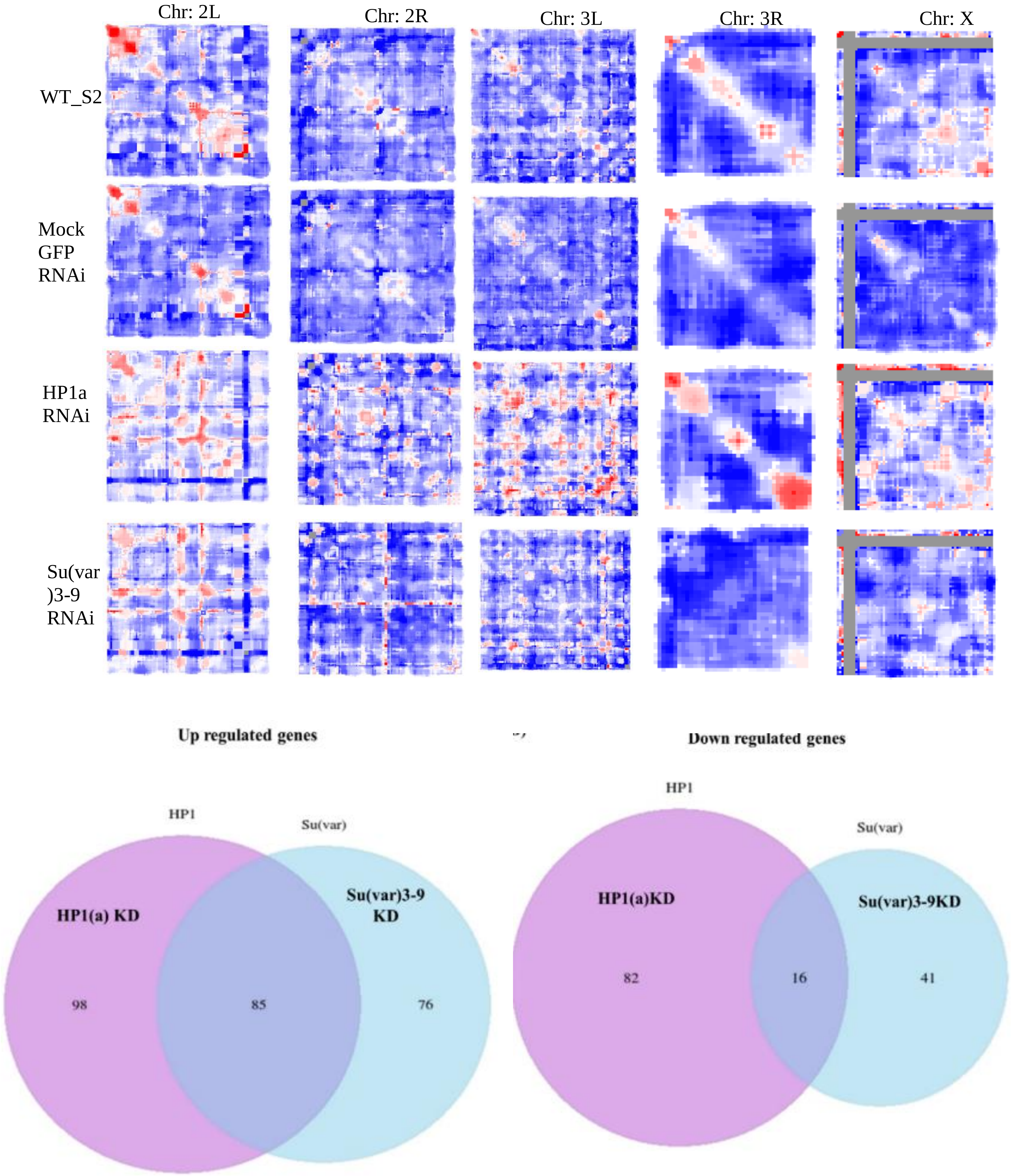


**Supplementary Figure S10: Effects of RNAi mediated depletion of HP1a or Su(var)3-9 on genome organization and gene expression A)** shows the heatmap for 5 chromosome arms in WT S2 cells, Mock GFP RNAi, HP1a RNAi and Su(var)3-9 RNAi. There is an increase in longer range DNA interactions not only across the diagonal but also on either side of it. **B)** Venn diagram showing the number of genes upregulated and down-regulated in HP1a and Su(var)3-9 knockdown conditions and the common between the two sets across the entire *Drosophila* transcriptome.

Table 2: List of differentially expressed genes in HP1a or Su(var)3-9 RNAi conditions ($p$ value < 0.05; Student two-sided T-test)

| **HP1a RNAi** | **Su(var)3-9 RNAi** |
|---|---|
| CG11739 | CG14636 |
| stnB | CG17159 |
| stnA | CG34357 |
| CG12581 | CG40006 |
| CG14615 | CG41434 |
| l(1)G0196 | CG6675 |
| Usp2 | CG9780 |
| CG14636 | CR33294 |
| CG17514 | CR41501 |
| Dsk | |
| Cht3 | |
| CG31525 | |
| Ir41a | |
| CG40006 | |
| CG40198 | |
| MFS17 | |
| Fog | |
| CG9780 | |
| CR41501 | |
| 18SrRNA-Psi:CR41602 | |
| Nipped B | |

**Table 3**: List of primers

| **Primers for validating 5C interactions by 3C-PCR** | |
|---|---|
| Primer name (chr –5Cprimer number) | Sequence (5’-3’) |
| 2L For8 | CCTGCCATCAATAGTCCGAAA |
| 2L For99 | AGCCGTATCGTCCGTAAG |
| 2L For 113 | CAGGTAACCATGAAAGCAAAGTAA |
| 2L For209 | GCTTAACAGCTCTCGCTCAT |
| 2R For139 | CCGACTTCCCACAGAACAAA |
| 2R For251 | GAGCACGAAAGCATCAGAATTG |
| 2R For259 | CCTTCGGAATGTCATCCTCATT |
| 2R For270 | GCCAGAAGAGAGTACTGACAAA |
| 2R For280 | GGCTGACACCAGTCAAGAAA |
| 2R For312 | CGTCTCCCACAAGAAGCTTAAA |
| 2R For317 | TATCGACCATACTGTCGAACTTT |
| 2R For423 | TACTTTGAGGGCACGCTTT |
| 3L For5 | GAGTGACAAGAAGCGTCTGC |
| 3L For36 | CACTCACCTCACGCAAT |
| 3L For61 | GAGGAACATACCCGGCATAC |
| 3L For164 | ACGACTTCTGGTAGCTCTCTAA |
| 3L For 441 | AGACGAGTCCATCGGAACTA |
| 3R For27 | ACTCCCAAGTGCACAGTAATC |
| 3R For62 | CCTTGTGCTGGTCACACTTAT |
| 3R For73 | CAGAGTGGTCAGGTACTTCTTC |
| 3R For85 | GCCTTAATTGTCCGCAGATT |
| X For67 | TTGTACTCCTCGATGAAGCTG |
| X For85 | GCCTTAATTGTCCGCAGATTG |
| X For189 | CCACTCGCCACCTCTTTG |
| X For215 | AAGATGGCGGTGTCCTTG |
| **Primers for FISH probes** | |
| 2LA1.F | ATCTACCTTGAACTGGGA |
| 2LA1.Rl | TGTGCTCAGGCGTCAGAAT |
| 2LA2.F | CAGGAAGACTGCTCAGGTAT |
| 2LA2.R | GGTCAGAACCATGCCGTAGT |
| 2LA3.F | GCTAGGCTCTAAGTATGAA |
| 2LA3.R | TCGCAGGTATCGGGTTGACA |
| 2LA4.F | ATAGGGCCAGGAGTGTTCGT |
| 2LA4.R | TGTGCAGGCATGTGCTTAAGTC |
| 2LB1.F | CGTCGGCTTGAGCTCGTCAT |
| 2LB1.R | GGCTGTGCACATTTGCT |
| 2LB2.F | ATGCCGTAAGCTTGTTGCG |
| 2LB2.R | AGTTTCGCACACGTCAACT |
| 2LB3.F | GTCAAGTATGTCTTTCCT |
| 2LB3.R | CTGGTGGCGTCATTGTCT |

| | |
|---|---|
| 2LB4.F | TGAGTGGCAAAGGTAACT |
| 2LB4.R | AAGGCAACAGGTGGGTGTAG |
| **Primers for RNAi mediated knockdown** | |
| EGFP_F | taatacgactcactatagggaCTACGGCGTGCAGTGCTTCA |
| EGFP_R | taatacgactcactatagggaCTACGGCGTGCAGTGCTTCA |
| HP1.F1 | taatacgactcactatagggaGCCCTCTGGCAATAAATCAA |
| HP1.R1 | taatacgactcactatagggaCAGGATAGGCGCTCTTCGTA |
| HP1.F2 | taatacgactcactatagggaTGGAGTACTATCTGAAATGGAAGG |
| HP1.R2 | taatacgactcactatagggaCGCCTCGTACTGCTGGATA |
| Su(var)3-9f | taatacgactcactatagggaTATTGAATGCGTCGAGATGG |
| Su(var)3-9r | taatacgactcactatagggaAGAGGTTCTCGTAGGGCACA |
| ADD1 F1 | taatacgactcactatagggaACCCCAACGTAGATCTGGTG |
| ADD1 F2 | taatacgactcactatagggaTGCTTGCGAGTATCTTGTGG |
| ADD1 R1 | taatacgactcactatagggaCAGAAGACGTAGGGGCAAGT |
| ADD1 R2 | taatacgactcactatagggaGCTCCATCTCGCGTAGGTAA |
| ADD1_RT_fwd | ACCCCAACGTAGATCTGGTG |
| ADD1_RT_rev | CAGAAGACGTAGGGGCAAGT |
| **Primers for quantitative PCR** | |
| Alpha-cat fwd | TGACAATACAGGGCAATGCG |
| Alpha-cat rvs | GATGTTATTAACGCAGCTA |
| Cta fwd | CAGGATGCTTATGATGCCA |
| Cta rvs | GCTCTTCTAATGCCACG |
| CG17510 fwd | GACCTACTCGCCCGGATAA |
| CG17510 rvs | TACACCATGACTCTTCG |
| CG17490 fwd | AGGGAGCGAACTTGAT |
| CG17490 rvs | AACTGGCTCGTACGCA |
| CG40298 fwd | GTGACTCCGAAGATATTG |
| CG40298 rvs | GGACTTGGCGTAGTTTA |
| nipped A fwd | GGAATATATATGGGTGGCGCAC |
| nippedA rvs | TGTCGCCACATCTCAGCTAT |
| p120ctn fwd | ACTGCTATACGTCGGATG |
| p120ctn rvrs | CTTCCAATCTTTGCGGCACT |
| light fwd | AGTCGAAGGCAGTGTT |
| light rvs | CAACGAAGGTGGCATCG |
| RpL15 fwd | TTCAGACTGCAGCTTGGC |
| RpLl5 rvs | ACCAGGCAATTCGTAACG |
| DIP1_fwd | GATCATCATCCGGTGCATCG |
| DIP_rvs | GATTCACGGCCGTTGTCAC |
| **3C controls** | |
| K1 | AAGCCGCAGGAGTTTCTAAC |
| K2 | CACGGGAAAACTACTGAAAG |

**Table 4:** List of antibodies

| Name | Source | Dilutions used |
|---|---|---|
| Anti HP1a | DSHB (C1A9) | 1:1000 |
| Anti Su(var)3-9 | Abcam ab4811 | 1:1000 |
| Anti Lamin | In-house (dm0) | 1:2000 |
| Anti Tubulin | Abcam ab6046 | 1:1000 |
| Anti Alexa fluor 488 | Molecular Probes | 1:400 |
| Anti Alexa fluor 555 | Jackson | 1:400 |
| Anti H3K9me3 | Abcam ab8898 | 2-3µg per ChIP reaction |
| Anti H3K36me3 | Abcam ab9050 | 2-3µg per ChIP reaction |

**Table 5**: List of Next Generation Sequencing datasets of S2 cell line used from NCBI GEO database or modENCODE

| **Genomic Feature** | **Experimental technique** | **Source** |
|---|---|---|
| H3K9me2 | ChIP seq | modENCODE_3953 |
| H3K9me3 | ChIP Chip | modENCODE_313 |
| H3K9ac | ChIP Chip | modENCODE_3765 |
| H3K27ac | ChIP Chip | modENCODE_3757 |
| H3K4me1 | ChIP Chip | modENCODE_3760 |
| H3K4me3 | ChIP Chip | modENCODE_3761 |
| H3K36me3 | ChIP seq | modENCODE_4715 |
| HP1a | ChIP Chip | modENCODE_3777 |
| HP1b | ChIP Chip | modENCODE_3020 |
| HP1c | ChIP Chip | modENCODE_3291 |
| mod(mdg4) | ChIP Chip | modENCODE_3789 |
| Su(Hw) | ChIP Chip | modENCODE_330 |
| CP190 | ChIP Chip | modENCODE_3748 |
| GAF | ChIP Chip | modENCODE_3753 |
| BEAF32 | ChIP on chip | modENCODE_3745 |
| CTCF | ChIP seq | modENCODE_2638 |
| dMES4 | ChIP seq | GSE56101 |
| dADD1 | ChIP seq | GSE56101 |
| Enhancers | STARR-Seq | GSE40739 |
| Boundary Elements | Bioinformatics prediction | cdBEST |
| MARs | Sequencing | Data from lab, available upon request |
| dMES4 KD | RNA seq | GSM1376614/15 |
